# Structural insights into ligand recognition and activation of the melanocortin-4 receptor

**DOI:** 10.1101/2021.06.21.449233

**Authors:** Huibing Zhang, Li-Nan Chen, Dehua Yang, Chunyou Mao, Qingya Shen, Wenbo Feng, Dan-Dan Shen, Antao Dai, Shanshan Xie, Yan Zhou, Jiao Qin, Jinpeng Sun, Daniel H. Scharf, Tingjun Hou, Tianhua Zhou, Ming-Wei Wang, Yan Zhang

**Author notes:** Correspondence (Y.Z.); (M.-W.W.). These authors contributed equally to this work (Huibing Zhang, Li-Nan Chen, Dehua Yang).

## Abstract

Melanocortin-4 receptor (MC4R) plays a central role in the regulation of energy homeostasis. Its high sequence similarity to other MC receptor family members, low agonist selectivity and the lack of structural information concerning receptor activation have hampered the development of MC4R-seletive therapeutics to treat obesity. Here, we report four high-resolution structures of full-length MC4R in complex with the heterotrimeric Gs protein stimulated by the endogenous peptide α-MSH, FDA-approved drugs afamelanotide (Scenesse™) and bremelanotide (Vyleesi™), and selective small-molecule ligand THIQ, respectively. Together with pharmacological studies, our results reveal the conserved binding mode of peptidic agonists, the distinctive molecular details of small-molecule agonist recognition underlying receptor subtype selectivity, and distinct activation mechanism for MC4R, thereby offering new insights into G protein coupling. Our work may facilitate the discovery of selective therapeutic agents targeting MC4R.

## INTRODUCTION

Melanocortin-4 receptor (MC4R), a key component of the leptin-melanocortin pathway, acts at the intersection of homeostatic maintenance of energetic state.^1-6^ Mutations in the MC4R gene were identified among patients with severe obesity from early childhood.^7, 8^ By far, natural occurring loss of function (LOF) MC4R mutations are the most frequent monogenic cause of obesity or binge eating disorder.^9^ MC4R agonists have been shown efficacious in appetite suppression, food intake reduction and body weight loss in people with high BMI (body mass index) caused by leptin, proopiomelanocortin (POMC) or MC4R deficiency.^10^ Obviously, MC4R plays a central role in regulating energy balance and satiety, and is one of the best validated drug targets for obesity.^11^ MC4R belongs to the melanocortin receptor family, a group of five (MC1R − MC5R) class A G protein-coupled receptors (GPCRs) and is primarily coupled to the stimulatory G protein (Gs) and increases intracellular cyclic adenosine monophosphate (cAMP) accumulation.^12^ MC1R governs mammalian skin and hair color by regulating the production of melanin.^13^ MC2R is mainly located in the adrenal cortex and controls the release of glucocorticoids.^2^ MC3R participates in the control of energy homeostasis and is implicated in immune responses, natriuresis and circadian rhythm.^14^ MC5R is involved in exocrine gland dysfunction, sebum regulation, and the pathogenesis of acne.^15–17^ Except MC2R, which is only activated by ACTH (adrenocorticotropic hormone) and requires an interplay with the melanocortin receptor accessory protein (MRAP) to attain functionality,^18^ the other four MCRs are endogenously activated by α-, β-, and γ-melanocyte-stimulating hormones (MSH), and blocked by the natural inverse agonist agouti-related peptide (AgRP).^19^ These features put forward a keen request for the development of a selective ligand for MC4R.

The endogenous ligands for MCRs contain a conserved His-Phe-Arg-Trp (HFRW) motif necessary for receptor activation (Supplementary information, Table S1).^20, 21^ Among them, α-MSH (Ac-Ser-Tyr-Ser-Met-Glu-His-Phe-Arg-Trp-Gly-Lys-Pro-Val-NH2) is the most well-studied peptide that has pleiotropic functions in pigmentation, cardiovascular system, erectile ability, energy homeostasis and immune responses.^21–24^ Driven by the pervasive issue of human obesity, tremendous efforts have been made to design potent and selective therapeutic agents targeting MC4R, both of peptidic and small-molecule nature.^23–27^ The linear peptide afamelanotide, Ac-Ser-Tyr-Ser-Nle^4^-Glu-His-D-Phe^7^-Arg-Trp-Gly-Lys-Pro-Val-NH2 (also known as [Nle^4^, *D*Phe^7^]α-MSH, NDP-α-MSH, melanotan-I, and MT-I), is capable of enhancing metabolic stability with markedly improved potency compared to α-MSH.^21, 28^ Subsequent efforts in obtaining peptides with better bioactivities led to the discovery of a cyclic peptide agonist, Ac-Nle^4^-c [Asp^5^, D-Phe^7^, Lys^10^] α-MSH(4-10)-NH2 (melanotan-II, MT-II).^21, 22^ Bremelanotide (C-terminal OH-MT-II) was approved in 2019 by the FDA to treat premenopausal women with sexual excitement dysfunction.^29^ Afamelanotide binds to MC1R in dermal cells to ramp up melanin production, effectively tanning the skin, and as a photo protectant received market approval in the European Union in 2014 and by the FDA in 2019.^30^

Unfortunately, owning to the high sequence homology among MCRs, most peptidic agonists initially developed for MC4R lack the capability of avoiding cross-reactivity with other MCR subtypes, such as MC1R, leading to adverse effects, *e.g.,* skin pigmentation and nausea.^10, 23, 29^ Intriguingly, subtype selectivity of MC4R was first achieved by a small-molecule agonist, THIQ, initially developed by Merck (1400-fold *vs*. MC1R, 1200-fold *vs*. MC3R and 360-fold *vs*. MC5R),^25^ that significantly decreases food consumption and simulates erectile responses.

The activation mechanism of MC4R and how it recognizes peptidic and small-molecule agonists are poorly understood, hampering the structure-guided drug discovery. Here, we employed single-particle cryo-electron microscopy (cryo-EM) to determine four structures of the human MC4R-Gs complexes: three bound to α-MSH, afamelanotide or bremelanotide at the resolutions of 3.0 Å, 3.0 Å and 3.1 Å, respectively; and one bound to THIQ at the resolution of 3.1 Å (Fig. 1; Supplementary information, Fig. S1, S2, Table S2). These structures reveal the molecular basis of ligand recognition and receptor activation for MC4R, shed light on the subtype selectivity of MCRs, and provide templates for the rational design of novel therapeutics targeting MC4R to treat obesity.

**Fig. 1.**
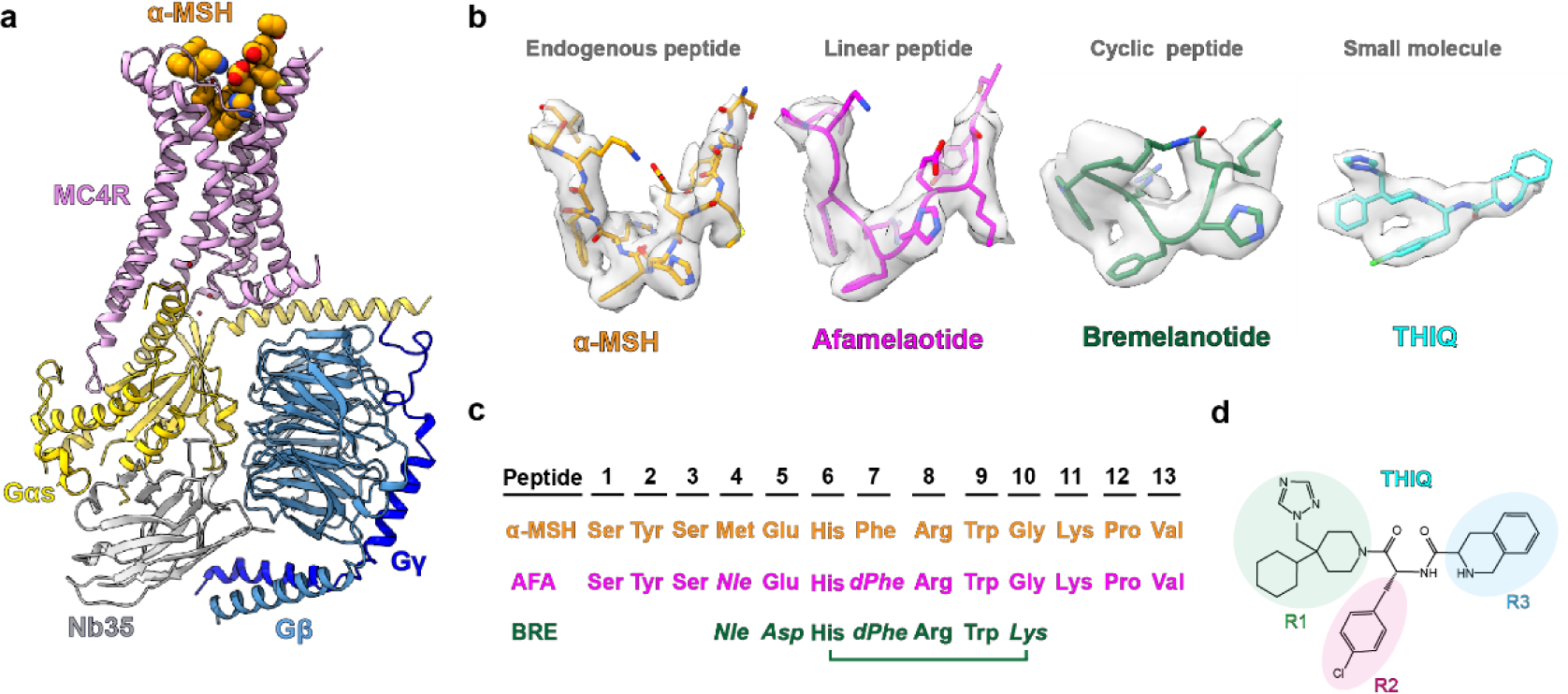
Cryo-EM structures of the MC4R–Gs complexes. **a** Cartoon representation of the α-MSH–bound MC4R–Gs complex. MC4R, plum; α-MSH, orange; Gαs, yellow; Gβ, cornflower blue; Gγ, blue; Nb35, grey. **b** Cryo-EM densities and models of α-MSH, peptide analogs (afamelanotide, bremelanotide) and small-molecule THIQ from their respective MC4R–Gs complexes. α-MSH, orange; afamelanotide, magenta; bremelanotide, forest green; THIQ, cyan, density is shown at 0.025 contour level. **c** Amino acid sequence of the agonists used in this study. Sequence alignment of three peptide agonists of MC4R (left). The mutant residues of afamelanotide (AFA) and bremelanotide (BRE) relative to α-MSH are shown in italics (right). **d** Two-dimensional chemical structure of THIQ.

## RESULTS

### Overall structures of agonist–MC4R–Gs complexes

To obtain a homogenous and stable MC4R–Gs complex, full-length MC4R was cloned into a pFastBac1 vector with a LgBiT and double maltose binding protein (MBP) affinity tag at the C-terminus. A dominant negative Gαs (DNGαs), Gβ1 and Gγ2 were co-expressed with MC4R in insect cells stabilized by a NanoBiT tethering strategy.^31–33^ Subsequently, assembly of the MC4R–Gs complex on the membrane was stimulated with an excess amount of agonist (α-MSH, bremelanotide, afamelanotide or THIQ (Fig. 1) and further stabilized by nanobody 35 (Nb35).^34^ The complex was then purified through affinity and size-exclusion chromatography to homogeneity, yielding a stable complex suitable for single-particle cryo-EM analysis (Supplementary information, Fig. S1a-b).

All four complexes, including α-MSH–MC4R–Gs–Nb35, afamelanotide–MC4R–Gs–Nb35, bremelanotide–MC4R–Gs–Nb35, and THIQ–MC4R–Gs–Nb35, were imaged under a Titan Krios microscope equipped with K2 summit direct detector, and the structures were determined at the global resolutions of 3.0 Å, 3.0 Å, 3.1 Å, and 3.1 Å, respectively (Supplementary information, Fig. S1c-f, S2a-d, Table S2). The high-resolution density maps showed that the majority of side chains in the signaling complexes were well resolved and allowed us to model most regions of MC4R from residues G39^N-term^ to Y320^8.61^ (superscript indicates the Ballesteros–Weinstein numbering system for class A GPCR)^35^ except G233^ICL3^ to R236^ICL3^ and D111^ECL1^ to Q115^ECL1^; α-MSH, afamelanotide, bremelanotide and THIQ, the Gs heterotrimer, and Nb35 in the final models were refined against the corresponding density map with excellent geometry (Fig. 1a-b; Supplementary information, Fig. S2). Additionally, a divalent calcium ion (Ca^2+^) was consistently observed in the peptide- and small-molecule-bound structures (Supplementary information, Fig. S2e-f, right, green sphere). Therefore, our structures could provide detailed information of the ligand binding mode between peptidic/small-molecule ligands and the receptor helix bundle, as well as the interface between Gs and MC4R, allowing us to examine conformational changes of the receptor upon agonist binding and the receptor-transducer coupling.

The overall conformations of MC4R–Gs complexes are almost identical in all four active structures (Cα root mean squared deviation (r.m.s.d.) < 0.3 Å), so we used the well-resolved α-MSH–MC4R–Gs structure (Fig. 1a) for further analysis below unless otherwise noted. Although MC4R adopts the canonical seven transmembrane (7TM) spanning helices architecture, it also displays several distinctive structural features. First, MC4R does not possess the conserved disulfide bond that usually connects the extracellular tip of TM3 to ECL2 in most GPCRs.^36^ However, the lack of a conserved disulfide bond also occurs in other class A GPCRs, such as sphingosine-1-phosphate receptors S1P1R-S1P5R, lysophosphatidic acid receptors LPA1R-LPA3R and cannabinoid receptors CB1R-CB2R.^37–42^ Second, three cysteine residues in ECL3 (C271, C277 and C279) are completely conserved among MCRs, two of which (C271 and C277) form an intra-ECL3 disulfide bond and C279 forms an additional disulfide bond with C40 in the N terminus (Supplementary information, Fig. S2e) which was not observed in the crystal structure of the antagonist SHU9119-bound thermostabilized MC4R (tsMC4R) in the inactive state.^43^ At the cytoplasmic side, the 7TM bundle of MC4R created a cavity to accommodate the heterotrimeric Gs protein. Unique to cryo-EM structures of MC4R–Gs complexes, several water molecules were trapped in the receptor–Gs interface (Fig. 1a). Notably, one water molecule unprecedentedly bridges R^1473.50^ of the conserved DRY motif with E392^G.H5.24^ (common Gα numbering system ^44^) in the C-terminal of Gαs (Fig. 1a).

Most class A aminergic, peptide and lipid GPCRs contain an elongated ECL2 segment consisting of over 15 amino acids and forming a β-sheet or helix lid, which usually covers the orthosteric binding pocket (Supplementary information, Fig. S3a-i). In contrast to these class A GPCRs, the ECL2 of MC4R is extremely short with only three amino acids, resulting in a widely open and extensively solvent-accessible orthosteric binding pocket.

### Recognition of α-MSH by MC4R

The extensive interactions between the endogenous agonist α-MSH and MC4R involve all seven TM helices plus the extended N terminus, ECL2 and ECL3 (Fig. 2a-b; details are presented in Supplementary information, Table S3). Electrostatic surface analysis revealed that the orthosteric binding pocket exhibits obvious amphiphilic characteristics, with the TM 2-3-4 half negatively charged while the rest of the pocket nearly hydrophobic (Supplementary information, Fig. S4a). The α-MSH inserts into the binding pocket in a charged complementary fashion (Supplementary information, Fig. S4a-c). The high-resolution density map showed an unambiguous density corresponding to a Ca^2+^ ion, coordinated by two backbone carbonyl oxygen atoms of Glu5 and Phe7 in α-MSH (peptide residues are labeled as three letters) and three negatively charged residues (E100^2.60^, D122^3.25^ and D126^3.29^) in MC4R (Fig. 2a). The side chain of Tyr2 and Arg8 located in close proximity to D122^3.25^, D126^3.29^ and I185^4.61^, and Arg8 hydrogen bonded to S188 in ECL2 (Fig. 2a-b; details are presented in Supplementary information, Table S3). In addition, F51^1.39^, T101^2.61^, I104^2.64^, F284^7.35^, N285^7.36^and L288^7.39^ prop up one supporting point His6, whereas I185^4.61^, I194^5.40^, L197^5.43^ and L265^6.55^ accommodate another supporting point Trp9 of the U-shaped α-MSH (Fig. 2a-b; details are presented in Supplementary information, Table S3). We mutated these residues that contact α-MSH to alanine and observed a significant loss of agonist-induced signal transduction in mutants F51^1.39^A, E100^2.60^A, D122^3.25^A, D126^3.29^A, I185^4.61^A, H264^6.54^A and L288^7.39^A, confirming the important roles of these residues in α-MSH–induced receptor activation (Fig. 2c; details are presented in Supplementary information, Table S3).

**Fig. 2.**
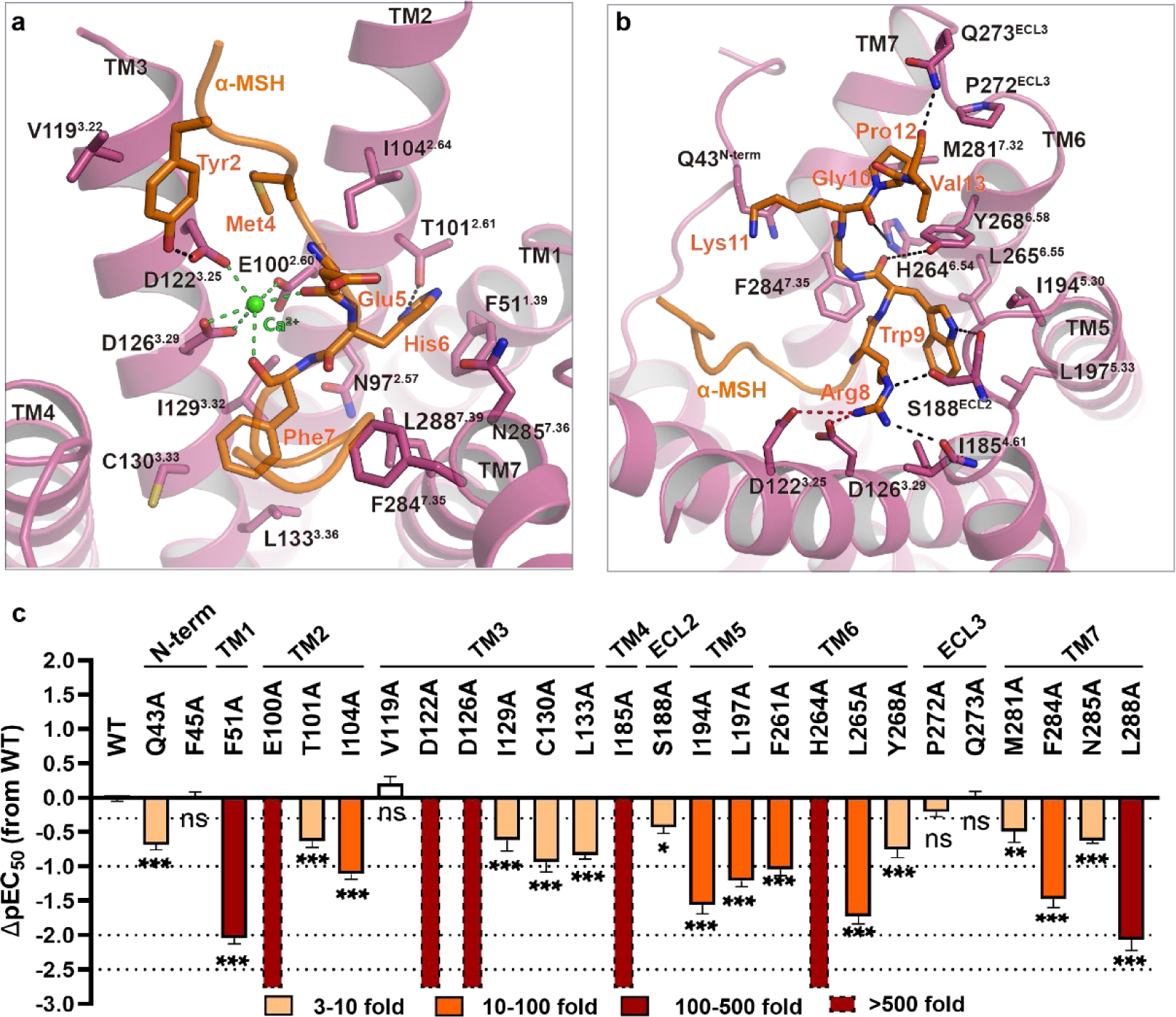
The binding pocket of α-MSH in MC4R. **a-b** Detailed interactions of α-MSH (orange) with MC4R (plum). Structure viewed from the extracellular side shows the interaction network between MC4R, α-MSH, and Ca^2+^. **c** α-MSH–induced cAMP accumulation assays of the residues involved in α-MSH binding. Bars represent differences in calculated α-MSH potency [pEC50] for representative mutants relative to the wild-type receptor (WT). Data are colored according to the extent of effect. \*\**P* > 0.01, \**P* < 0.01, \*\**P* < 0.001 and \*\*\**P* < 0.0001 (one-way ANOVA followed by Dunnett’s multiple comparisons test, compared with the response of WT). See Supplementary Table S4 for detailed statistical evaluation and receptor expression levels.

The EM density map indicated that Glu5 of α-MSH possibly adopted two distinct rotamers, inducing local changes of the α-MSH–MC4R interface environment (Supplementary information, Fig. S4g-h). One rotamer of Glu5 side chain is likely to direct towards Lys11 forming a polar interaction, linking two flanks of the peptide. The resulting cyclized ring-like conformation resembles the conformation of cyclic peptide agonist bremelanotide (Fig. 3a, b). The other rotamer of Glu5 points down towards His6. By adopting a similar direction, the aspartate at the corresponding site (Asp3) of SHU9119 formed a hydrogen bond with His6 in the crystal structure of tsMC4R.^31^ In this case, His6 of α-MSH hydrogen bonded to N285^7.36^ and T101^2.61^ (Supplementary information, Fig. S4g-h). Notably, tsMC4R contained a substitution of N97^2.57^L, which folded towards the bottom of the ligand biding pocket. Interestingly, we observed the side chain of N97^2.57^ in the α-MSH–MC4R structure adopting a conformation similar to that in the tsMC4R structure (Supplementary information, Fig. S4g-h). In agreement with the critical role of N97^2.57^ in receptor activation, substitution of N97^2.57^ with leucine or alanine resulted in partial (about 25%) or complete loss of α-MSH–induced cAMP responses (Supplementary information, Fig. S4i; details are presented in Supplementary information, Table S4). N97^2.57^A mutation resulted in near-complete loss of surface trafficking (Supplementary information, Table S4), and consequent dramatic reduction in activity (Supplementary information, Table S5) and a corresponding cAMP response (Supplementary information, Fig. S4i). However, although the mutant N97^2.57^L maintained the robust surface expression level (Supplementary information, Table S4) and binding affinity of α-MSH (Supplementary information, Table S5), it partially lost cAMP response (Supplementary information, Fig. S4i).

**Fig. 3.**
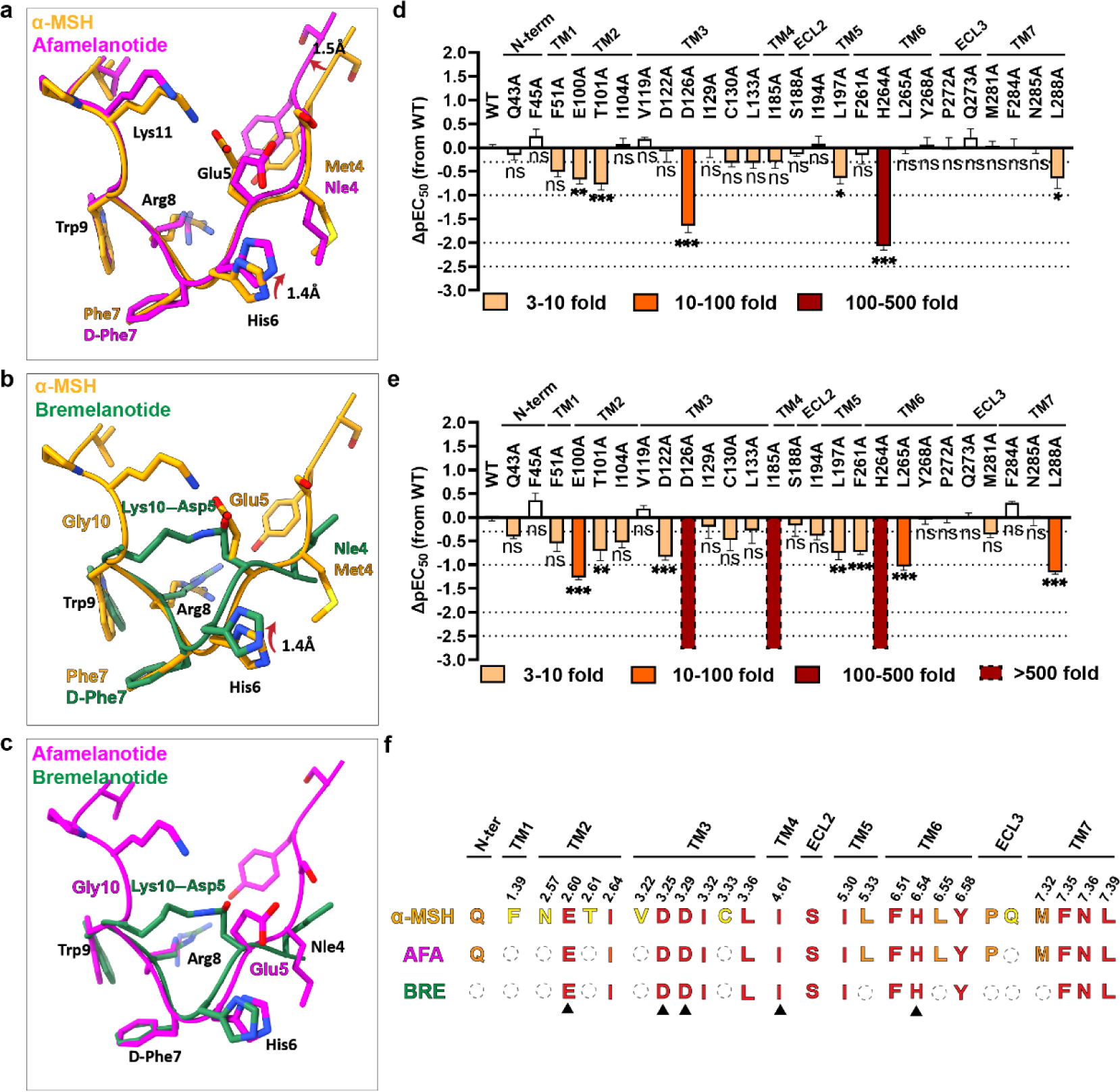
Differences in the recognition mode of three peptide agonists of MC4R. **a-c** Parallel structural comparisons of α-MSH–afamelanotide, α-MSH–bremelanotide and afamelanotide–bremelanotide. The peptide ligands are superimposed upon alignment of MC4R. α-MSH, orange; afamelanotide, magenta; bremelanotide, forest green. **d-e** Afamelanotide-(**d**) and bremelanotide-induced cAMP accumulation (**e**) assays of the residues involved in ligand recognition. Bars represent differences in calculated agonist potency [pEC50] for each mutant relative to the wild-type receptor (WT). Data are colored according to the extent of effect. **^ns^***P* > 0.01, \**P* < 0.01, \*\**P* < 0.001 and \*\*\**P* < 0.0001 (one-way ANOVA followed by Dunnett’s multiple comparisons test, compared with the response of WT). See Supplementary Table S7 and S8 for detailed statistical evaluation and receptor expression levels. **f** Comparison of the binding site of α-MSH, afamelanotide (AFA) and bremelanotide (BRE) in MC4R. The triangle labeled residues are crucial for MC4R activation.

### Recognition of afamelanotide and bremelanotide by MC4R

Sole replacement of Phe7 in α-MSH with D-Phe7 produced afamelanotide with significant increase in binding affinity and functional activity.^21^ The recognition mode of afamelanotide by MC4R therefore is very similar to that of α-MSH (Fig. 3a, f; Supplementary information, Fig. S5a-c). Bremelanotide has less interactions with MC4R only involving TMs 2-7, owing to the omission of flanking residues at C and N termini (Fig. 3b, c; Supplementary information, Fig. S5d; details are shown in Supplementary information, Table S6, 7). Structural superimposition revealed that, although substitution of Phe7 with D-Phe7 did not alter the overall shape of afamelanotide compared to α-MSH, it indeed induced notable conformational changes in the peptide. The N-terminal flank of afamelanotide shifted 1.5 Å away from TMs 2-3 measured at the Cα of Ser1, resulting in the partial loss of interactions with TM2 and TM3 (Fig. 3a, f; Supplementary information, Fig. S5b, c). D-Phe7 almost stayed at the same position as Phe7 of α-MSH. However, the replacement led to the neighboring residue His6 swinging upward 1.4 Å to form a hydrogen bond with Glu5 (Fig. 3a), together with the breaking of the hydrogen bond with N97^2.57^ and π–π stacking with F51^1.39^ (Fig. 2a Supplementary information, Fig. S5b, c). Similar conformational changes of Glu5 and His6 were also observed in the structure of bremelanotide-bound MC4R (Fig. 3b, c, 3f; Supplementary information, Fig. S5d). Consistent with our observation that F51^1.39^ was not involved in the recognition of afamelanotide or bremelanotide, the cAMP assay showed that the substitution of F51^1.39^A only reduced afamelanotide- and bremelanotide-induced responses by between 3- and 10-fold in contrast to over 100-fold for α-MSH (Fig. 3d, e, details are shown in Supplementary information, Table S8, 9). Additionally, the distance between Glu5 and His6 in afamelanotide (3.4 Å) was in the preferable range of hydrogen bond compared to 4.6 Å in α-MSH (Fig. 3a). The more stabilized interaction between Glu5 and His6 in afamelanotide trapping the side chain of Glu5 points toward His6, rigidifying the local conformation.

### Specific Recognition of THIQ by MC4R

The cryo-EM map enabled the unambiguous assignment of tri-branched THIQ sitting in the bottom of the orthosteric binding pocket (Fig. 1b, d; Supplementary information, Fig. S5a). The binding of THIQ is mediated by residues from six transmembrane helices (TMs 2-7) as well as residues from the N terminus and ECL2 (Fig. 4a; details are given in Supplementary information, Table S10). Compared to the peptidic agonists discussed above, the conformational architecture of THIQ reassembled the His-Phe-Arg-Trp pharmacophore of α-MSH. Moiety R1 took the place of Trp9 of the peptide agonist (Fig. 4a, b). The cyclohexane group from R1 fitted to a hydrophobic pocket consisting of S188^ECL2^, I185^4.61^, I194^5.40^ and L197^5.43^, whereas triazole from R1 formed a tight hydrogen bond with H264^6.54^ and Van der Waals interactions with I194^5.40^, L265^6.55^ and Y268^6.58^ (Fig. 4a, b). Consistently, substitution of H264^6.54^ with alanine abolished the THIQ-induced activation (Fig. 4c). Moiety R2 D-Phe (*p*Cl) structurally mimicked Phe7, pointing to R1 and inserting into a hydrophobic pocket comprising I129^3.32^, C130^3.33^, L133^3.36^, F261^6.51^ and F284^7.35^. Moiety R3 plugged into another hydrophobic pocket in which His6 of peptidic agonists was accommodated by F45^N-term^, I104^2.64^ and L288^7.39^, and was furthermore in close proximity to N285^7.36^(Fig. 4a, b). However, similar to afamelanotide and bremelanotide, R3 of THIQ did not form a hydrogen bond with N285^7.36^ (Fig. 4a, b), whose replacement with alanine therefore did not affect the potency of THIQ (Fig. 4c; details are given in Supplementary information, Table S11). In contrast with peptidic agonists, only one carbonyl oxygen atom from THIQ, equivalent to that of Phe7 in endogenous peptide ligand, participated in coordinating Ca^2+^ (Fig. 4a). Hence, it appears that the function of THIQ is less dependent on the metal ion when compared to α-MSH, as shown by our cAMP assay (Supplementary information, Fig. S5e, h; details are given in Supplementary information, Table S12).

**Fig. 4.**
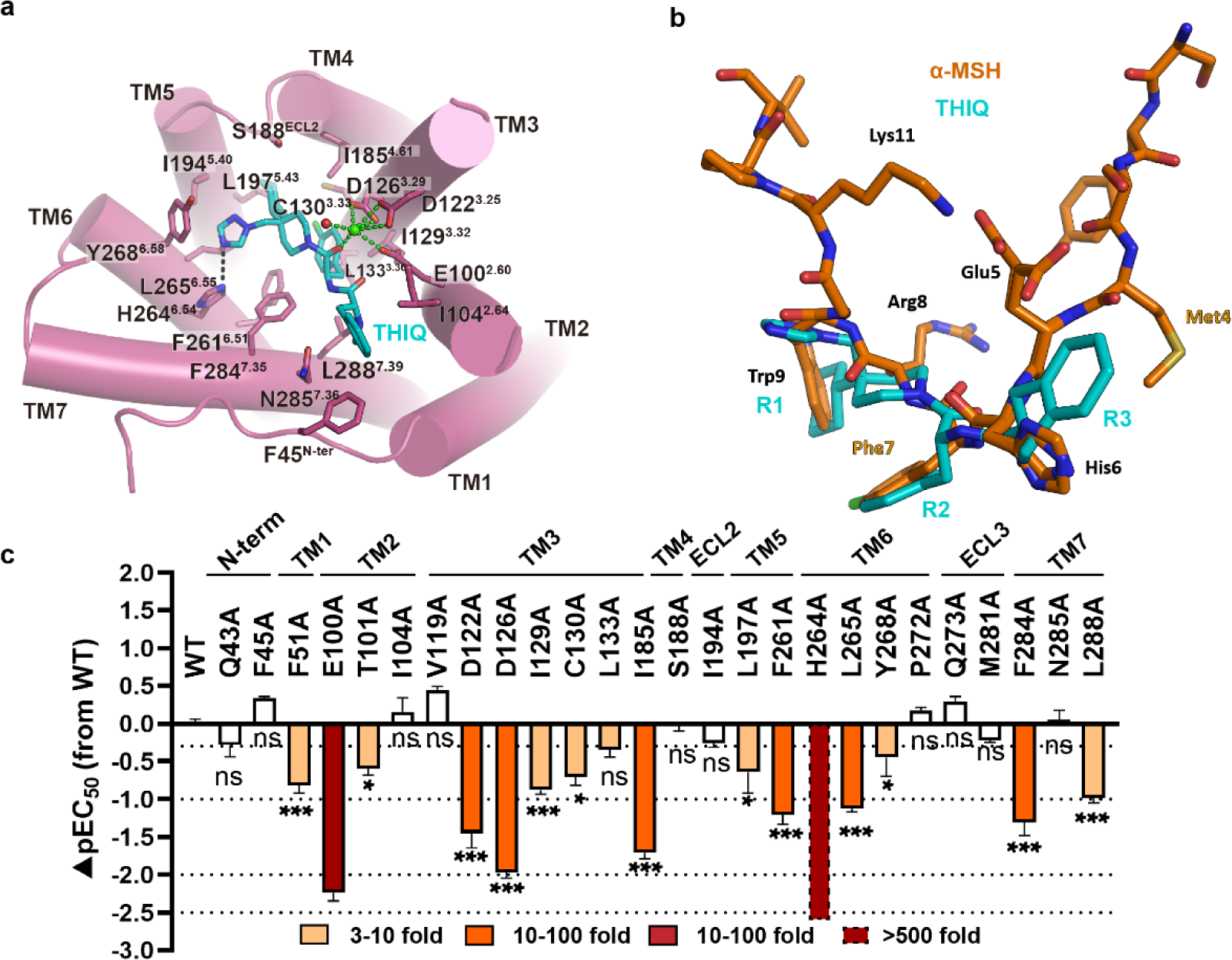
Interaction between THIQ and MC4R. **a** Detailed interaction of small-molecule THIQ (cyan) with MC4R (plum). Structure viewed from the extracellular side shows the interaction network between MC4R, THIQ and Ca^2+^. **b** Structure comparison of α-MSH with THIQ. The two ligands are aligned by the receptor. **c** THIQ-induced cAMP accumulation assays of the residues involved in THIQ binding. Bars represent differences in calculated α-MSH potency [pEC50] for each mutant relative to the wild-type receptor (WT). Data are colored according to the extent of effect. **^ns^***P* > 0.01, \**P* < 0.01, \*\**P* < 0.001 and \*\*\**P* < 0.0001 (one-way ANOVA followed by Dunnett’s multiple comparisons test, compared with the response of WT). See Supplementary Table S10 and S11 for detailed statistical evaluation and receptor expression levels.

In addition, we also docked several other small-molecule agonists into MC4R to characterize the general activation mechanism of MC4R induced by non-peptidic compounds. The docking poses of CHEMBL406764, MB-243 and RY764 highly resembled the binding pose of THIQ, in which the p-chlorophenyl group of these compounds inserted into the same hydrophobic pocket of MC4R as THIQ (Supplementary information, Fig. S6a-f). Particularly, all of these three compounds shared the same Ca^2+^ coordination, consistent with our observation of the THIQ-MC4R complex structure (Supplementary information, Fig. S5d). However, the docking poses indicate that moiety R3 was more flexible compared to moieties R1 and R2, in which CHEMBL406764 is nearly perpendicular to THIQ at moiety R3 (Supplementary information, Fig. S6d-f). This is probably due to the fact that the pockets for moieties R1 and R2 are much deeper and more specific so that THIQ has relatively more extensive interactions in these two pockets, whereas the pocket for moiety R3 is shallow and flat in which THIQ has nonspecific hydrophobic interactions with only three residues*, i.e.*, F45^N-term^, I104^2.64^ and L288^7.39^. Nevertheless, the structure of the THIQ-MC4R complex provides a template to understand the structure-activity relationship (SAR) of small molecules targeting MC4R and facilitates the rational design of drugs with the desired activity.

### Receptor subtype selectivity

Structure-based sequence alignment suggests most of the residues in the peptidic ligand binding pocket are highly conserved among MCR family members. Moreover, critical residues for ligand binding and functional activity are completely conserved, such as E100^2.60^, D122^3.25^, D126^3.29^, I185^4.61^, F261^6.51^, H264^6.54^, L265^6.55^, F284^7.35^ and L288^7.39^ (Supplementary information, Fig. S7a), providing the molecular basis of polypharmacology of peptidic agonists of MCRs. Among the residues involved in selective THIQ recognition, three are non-conserved between MC1R and MC4R, *i.e.*, I129^3.32^ at TM3, S188 at ECL2 and Y268^6.58^ at the extracellular end of TM6, whereas T124^3.32^, Y183^ECL2^ and I264^6.58^ are present in the corresponding positions in MC1R (Fig. 5a, d). I129^3.32^ is involved in forming the hydrophobic pocket accommodating R2 group of THIQ, whereas Y268^6.58^ may interact with the imidazole ring. The substitution of S188^ECL2^ with bulkier residues may hamper the entry of THIQ. We thus anticipate that these three residues collaboratively determine the receptor subtype selectivity of THIQ. To test our hypothesis, we replaced them with the corresponding residues of MC1R and evaluated the potency of THIQ and α-MSH using the cAMP assay. It was found that substitution of non-conserved residues dramatically reduced the potency of THIQ by 5-20-fold (Fig. 5b; details are shown in Supplementary information, Table S11). The combination of S188^ECL2^Y and Y268^6.58^I mutations also significantly decreased the potency of THIQ over 50-fold (Fig. 5b, details are shown in Supplementary information, Table S11). Furthermore, when we replaced all three residues of MC4R with those of MC1R, THIQ-induced cAMP production was nearly eliminated (Fig. 5b, details are shown in Supplementary information, Table S11). In contrast, different mutation combinations resulted in negligible to mild effects on the functional activity of α-MSH (Fig. 5c; details are shown in Supplementary information, Table S4). This data shows that I129^3.32^, S188^ECL2^ and Y268^6.58^ in MC4R are critical residues responsible for selective recognition of THIQ by this receptor subtype.

**Fig. 5.**
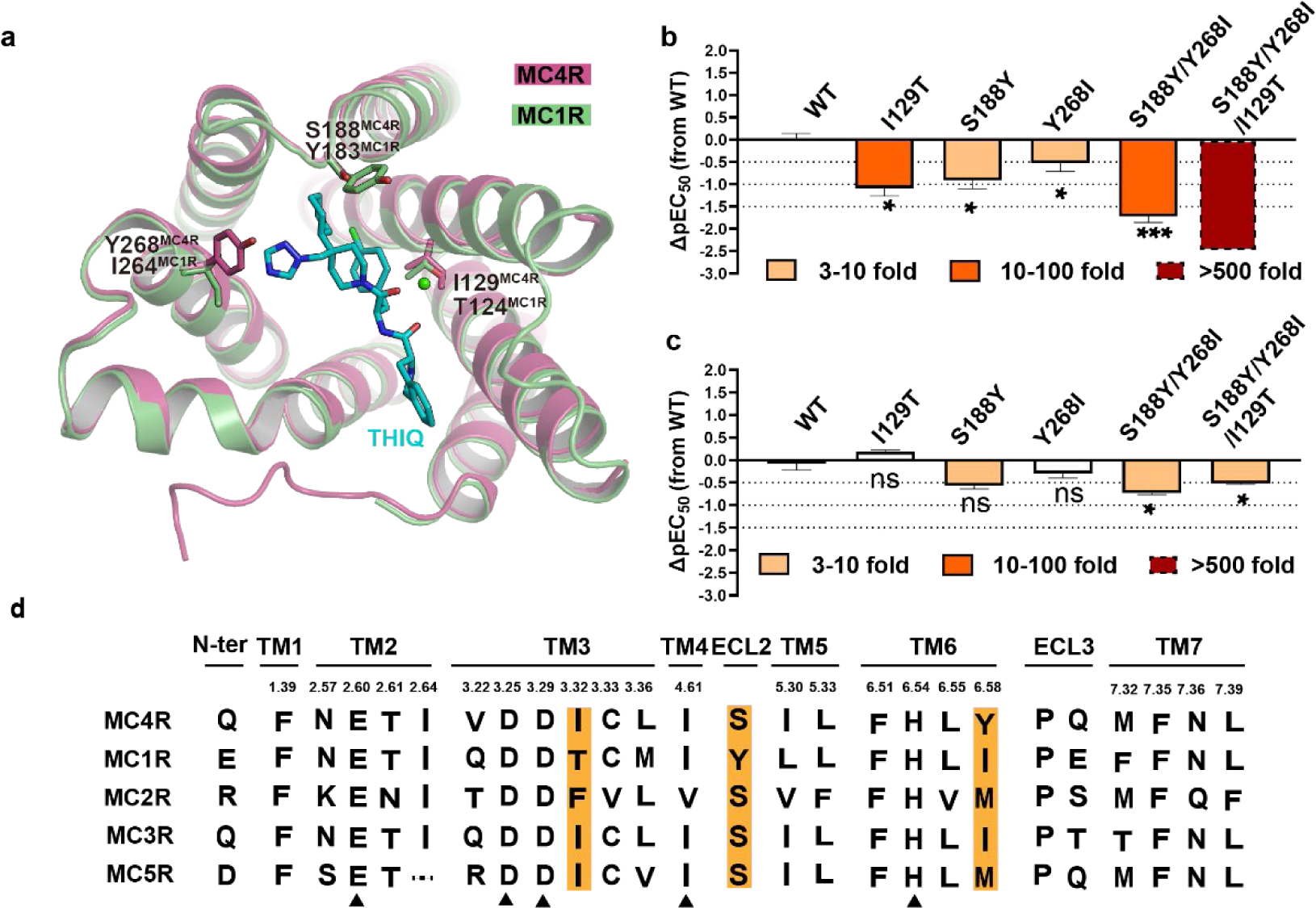
Selective activation of MC4R by THIQ. **a** Structure comparison of MC4R and the homology modeled structure of MC1R. The non-conserved residues in the THIQ binding pockets between the two receptors are shown as sticks. MC4R, plum; MC1R, green; THIQ, cyan. **b-c** THIQ– (**b**) and α-MSH–induced cAMP accumulation (**c**) assays of the potential residues involved in selectivity. Bars represent differences in calculated α-MSH potency [pEC50] for representative mutants relative to the wild-type receptor (WT). \*\**P* > 0.01, \**P* < 0.01, \*\**P* < 0.001 and \*\*\**P* < 0.0001 (one-way ANOVA followed by Dunnett’s multiple comparisons test, compared with the response of WT). See Supplementary Table S4 and S11 for detailed statistical evaluation and receptor expression levels. **d** Sequence alignment of the residues in the orthostatic binding pocket of MC4R and other MCR family members. Highlighted are the orthostatic binding pocket residues that contribute to selectivity.

In summary, the sequence alignment of MC4R and MC1R displays 19 residue variants in the orthosteric binding pocket, including 15 residues exhibiting different chemical properties and 4 residues with similar properties (Supplementary information, Fig. S7b-d). Structurally, most variants are located at the periphery of the 7TM extracellular side. However, we noticed that two variants, I129^3.32^ and L133^3.36^ (T124^3.32^ and M128^3.36^ in MC1R) reside at the bottom of the ligand binding pocket, where they are in close proximity to the His-Phe-Arg-Trp pharmacophore. It is also noteworthy that four variants in the N terminus (Q43, L44, F45 and V46), one in TM6 (Y268^6.58^) and three in ECL3 (V278, M281 and S282) are located close to each other in a restricted region, making it a hotspot for ligand specificity (Supplementary information, Fig. S7b-d).

### Role of Ca2+ in ligand activity

Ca^2+^ is considered a necessary cofactor for ligand binding and activity.^43^ To evaluate the effect of Ca^2+^ on the recognition of different ligands by MC4R, we gradually added EGTA to remove Ca^2+^ from the medium and evaluated the binding and potency of the MC4R ligands in question (Supplementary information, Fig. S5e-h; details are given in Supplementary information, Table S12). As expected, α-MSH exhibited a complete dependency on Ca^2+^, evinced by the fact that the supplement of 5 mM EGTA abolished the activity of α-MSH (Supplementary information, Fig. S5e; details are given in Supplementary information, Table S12). The cyclic peptide bremelanotide behaved similarly in lacking cAMP production following the addition of EGTA at different concentrations (Supplementary information, Fig. S5g; details are given in Supplementary information, Table S9, 12). Surprisingly, afamelanotide and THIQ preserved 80% and 40% maximal efficacy of ligand-induced cAMP signaling in the presence of 5 mM EGTA, respectively (Supplementary information, Fig. S5f, h; details are given in Supplementary information, Table S12). Of the three MC4R residues involved in Ca^2+^ action (E100^2.60^, D122^3.25^ and D126^3.29^), mutating any of them would abolish the effect of α-MSH (Fig. 2a, c; details are given in Supplementary information, Table S4). Substitution of D126^3.29^A abolished bremelanotide-stimulated cAMP accumulation, but only decreased the activities of afamelanotide and THIQ about 40- and 100-fold, respectively. Similarly, mutant E100^2.60^A reduced the potencies of bremelanotide and THIQ about 170- and 20-fold, respectively, but had almost no effect on afamelanotide-induced MC4R activation (Fig. 3d, e, 4c; details are given in Supplementary information, Table S4, 8, 9, 12). It thus appears that the activity of these ligands is dependent on Ca^2+^ to varying degrees, raising the question of whether or not there is a Ca^2+^-independent mechanism for MC4R modulation.

### Activation mechanism of MC4R

In comparison with the recently reported structure of SHU9119-bound tsMC4R in the inactive state (PDB: 6W25),^43^ binding to an agonist did not induce notable conformational changes at the extracellular side of MC4R (Fig. 6a-b). This finding is in contrast to previous structural studies of class A GPCRs, in which agonist-induced receptor activation requires the extensive contraction of the 7TM extracellular half correlated with the opening of the receptor cytoplasmic side to create a cavity for G protein coupling.^36^ Therefore, the most profound conformational changes of MC4R upon activation were observed in the cytoplasmic regions that were stabilized by Gs engagement, including (i) TM6 intracellular end exhibited 9.8 Å outward movement measured at Cα of K242^6.32^; and (ii) 6.9 Å inward movement of TM5 measured at Cα of L217^5.63^ (Fig. 6a-e).

**Fig. 6.**
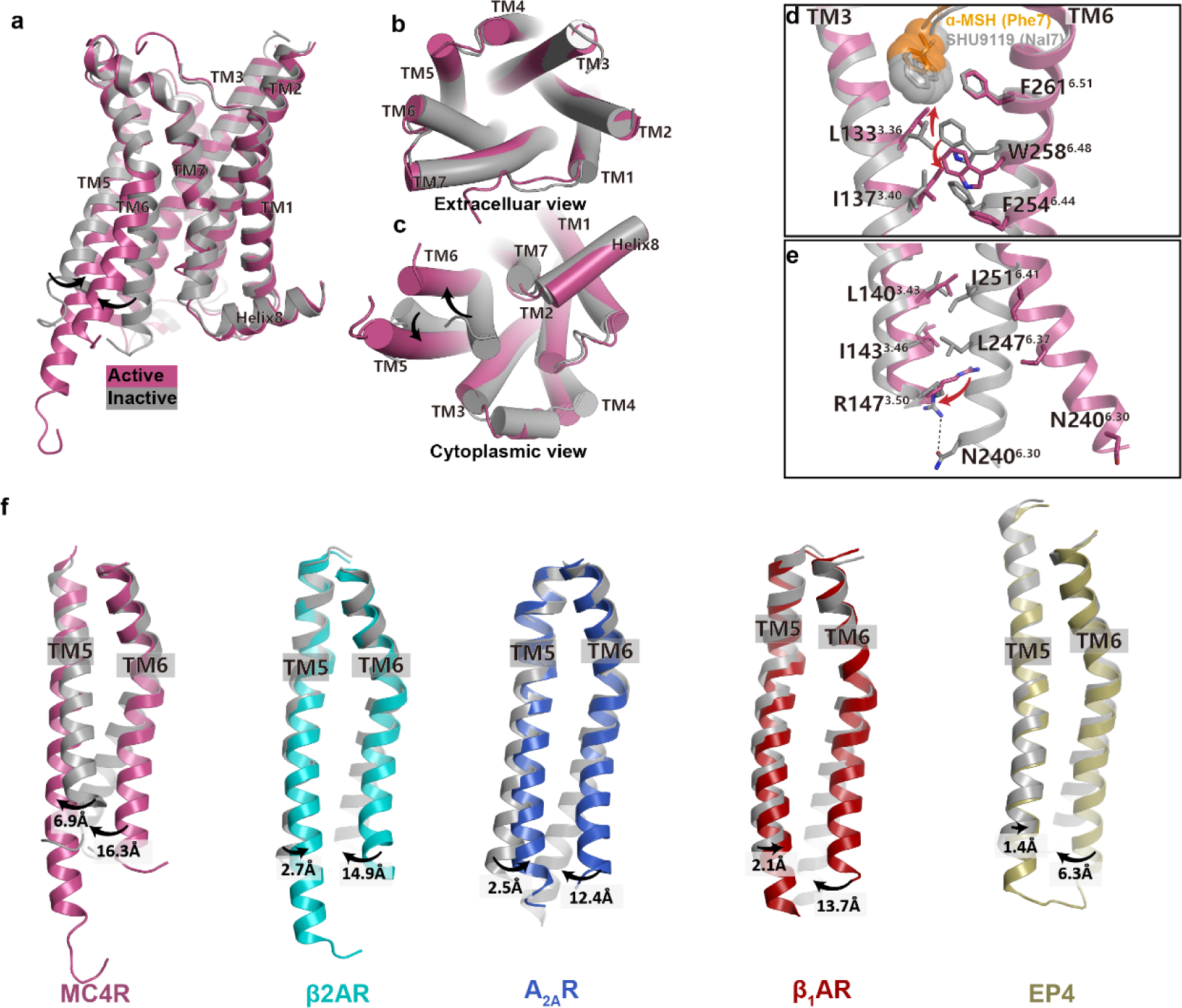
Activation mechanism of MC4R. **a-c** Superposition of the α-MSH-activated MC4R (plum) with the antagonist (SHU9119) –bound MC4R (grey) (PDB: 6w25). (**a**) Side view. (**b**) Extracellular view. (**c**) Cytoplasmic view. **d-e** Close-up views of the conformational changes of the crucial residues involved in MC4R activation. **f** Structure comparisons of TM5 movements among peptide receptors during activation. Inactive and active MC4R, grey and plum; inactive and active β2AR, grey (PDB: 6PS6) and cyan (PDB: 3SN6); inactive and active A2AR, grey (PDB: 6LPJ) and blue (PDB: 6GDG); inactive and active β1AR, grey (PDB: 3ZPQ) and red (PDB: 7JJO); inactive and active EP4, grey (PDB: 5YWY) and yellow (PDB: 7D7M).

Displacement of D-Phe7 in bremelanotide by the bulkier hydrophobic group D-naphthylphenylalanine (D-Nal) led to SHU9119 and this subtle change sufficiently turned a potent MC4R agonist into an antagonist (Fig. 6d; Supplementary information, Fig. S8a-f, Table S2).^23^ Therefore, peptidic agonists including α-MSH and SHU9119 shared a similar binding mode with MC4R, providing an explanation for the observation that negligible conformational changes at the extracellular side were observed upon receptor activation. The most profound conformational changes among the ligand interacting residues occurred at the position L133^3.36^. L/D-Phe7 of the agonists is 2 Å shorter than that of D-Nal7 in SHU9119 when inserted into the binding pocket, leading to the upward movement of L133^3.36^ followed by a 2.9 Å downward movement of toggle switch W258^6.48^ (Fig. 6d). Substitution of L133^3.36^ with alanine resulted in a dramatic decrease in basal activity of the receptor, highlighting the important role of L133^3.36^A as a sensor defining the nature of the ligand during receptor activation (Supplementary information, Fig. S8g). To support this hypothesis, we examined the SAR of peptide agonist derivatives with different modifications of D-Phe7.^23, 45^ Apparently, D-Phe7, D-Phe7(*p*F, *p* indicating a substituent in the *para* position of phenyl group) and D-Phe7(*p*Cl) in either peptidic or small-molecule agonists displayed a similar optimal potency, whereas D-Phe7(*p*I) caused a considerable steric hindrance with L133^3.36^ due to its larger Van der Walls radius reducing the efficacy by 33 folds compared to D-Phe7, D-Phe7(*p*F) and D-Phe7(*p*Cl) (Supplementary information, Fig. S8a-f). The bulky D-Nal group created a severe steric clash with L133^3.36^ in the active state, and thus pushed the side chain of L133^3.36^ to rotate downwards facing the cytoplasmic side accompanied by an upward movement of W258^6.48^, resulting in the “toggle twin switch” (L133^3.36^ and W258^6.48^) in a canonical inactive conformation of class A GPCR (Fig. 6d). On the other hand, swapping NDP-MSH residue D-Phe7 to D-Ala7 would result in the loss of the hydrophobic interaction between the phenyl group and L133^3.36^, thereby insufficiently stabilizing L133^3.36^ in the upward active conformation (Supplementary information, Fig. S8a). Consequently, this substitution profoundly decreased ligand potency by over 1000-fold.^39^ Markedly, substituting the residue L133^3.36^ to methionine led to the complete conversion of SHU9119 activity from antagonist to agonist of MC4R,^46^ possibly owing to the fact that the methionine side chain has more freedom to adopt different rotamers accommodating the bulky D-Nal7 residue of SHU9119, further emphasizing the critical role of L133^3.36^ in receptor activation. In conclusion, our agonist-bound structures uncovered the MC4R agonism which is precisely modulated by the spatial size of the residue located at the peptide 7-position or the equivalent moiety of small-molecule compounds (Supplementary information, Fig. S8a-f).

The concerted conformational changes of the “toggle twin switch” L133^3.36^ and W258^6.48^ relayed by M204^5.50^I137^3.40^F254^6.44^ motif in the MC receptor family, are equivalent to the conserved PIF motif in other class A GPCRs.^47^ The above-mentioned agonist-induced structural changes drove W258^6.48^ to slide downward contacting L133^3.36^ and I137^3.40^ propagated through F254^6.44^ (Fig. 6d). The outward movement of F254^6.44^ opened up TM6 by breaking the hydrophobic interaction between L140^3.43^ and I143^3.46^ in TM3 and I251^6.41^ and L247^6.37^ in TM6 and the hydrogen bond between R147^3.50^ and N240^6.30^, creating a cytoplasmic cavity for α5 of Gαs (Fig. 6e). This suggests that F254^6.44^ also plays a key role in MC4R activation

Interestingly, TM5 exhibited a profound yet unique conformational change induced by agonist and G protein binding (Fig. 6f). The structure superposition showed that TM5 extended three helical turns and moved inward 6.9 Å measured at the Cα atom of L217^5.63^ upon activation (Fig. 6a, f). The lipid separating TM3 and TM5 in the inactive MC4R was squeezed off from the activated receptor as we did not observe any density at the equivalent position (Supplementary information, Fig. S8h-j). The interactions between TM3 and TM5 were fastened by the newly established hydrogen bond between Y148^3.51^ and H214^5.60^. The hydrogen bond between H214^5.60^ and H222^5.68^ in the inactive state was broken by the extension of TM5 (Supplementary information, Fig. S8h-j). In contrast, previous structural studies of class A GPCRs, such as β2AR, A2AR, β1AR and EP4, demonstrated that receptor activation always induces an outward movement of TM5 to various extents, 1.4-2.8 Å measured at Cα of the residue 5.66 (β2AR: K277; A2AR: R205; β1AR: K235; and EP4: L211) (Fig. 6f).^48–55^ The distinctive structural rearrangement of TM5 related to MC4R activation probably attributes to the presence of a lipid between TM3 and TM5 in addition to the absence of the conserved Pro at 5.50.^47^

### MC4R-Gs coupling interface

The MC4R-Gs interface involves TM3, TMs 5-7 and all three ICLs (Fig. 7; details are shown in Supplementary information, Table S13). The overall cryo-EM structures of the MC4R-Gs complexes revealed a similar interaction mode of the MC4R-G protein compared to other class A GPCRs, such as β2AR-Gs (PDB: 3SN6), A2AR-Gs (PDB: 6GDG) and GPBAR-Gs (PDB: 7CFM) ^34, 49, 56^ (Supplementary information, Fig. S9a-b). However, the high-resolution density map showed several distinctive features of the MC4R-Gs complex. In other reported class A GPCR-Gs complexes, the side chain of Y391^G.H5.23^ in Gα helix 5 was recognized primarily by packing against TM3 in parallel with R^3.50^ from the conserved DRY motif with the distance of 4 Å between two side chain planes and did not established strong interactions with the receptors. In the MC4R-Gs structures, however, the hydroxyl group of Y391^G.H5.23^ formed a tight hydrogen bond with the side chain of T150^3.53^, a phenomenon strictly conserved in the MCR family. Residues at the equivalent position in other class A GPCRs with available structures in complex with Gs are alanine (Fig. 7f; Supplementary information, Fig. S9c-d). Strikingly, displacement of T150^3.53^ with alanine completely abolished the agonist-induced cAMP production but did not affect the β-arrestin recruitment to MC4R, suggesting that T150^3.53^ plays an important role in Gs coupling to MC4R but may not be involved in β-arrestin signaling (Fig. 7g-h; details are shown in Supplementary information, Table S14).

**Fig. 7.**
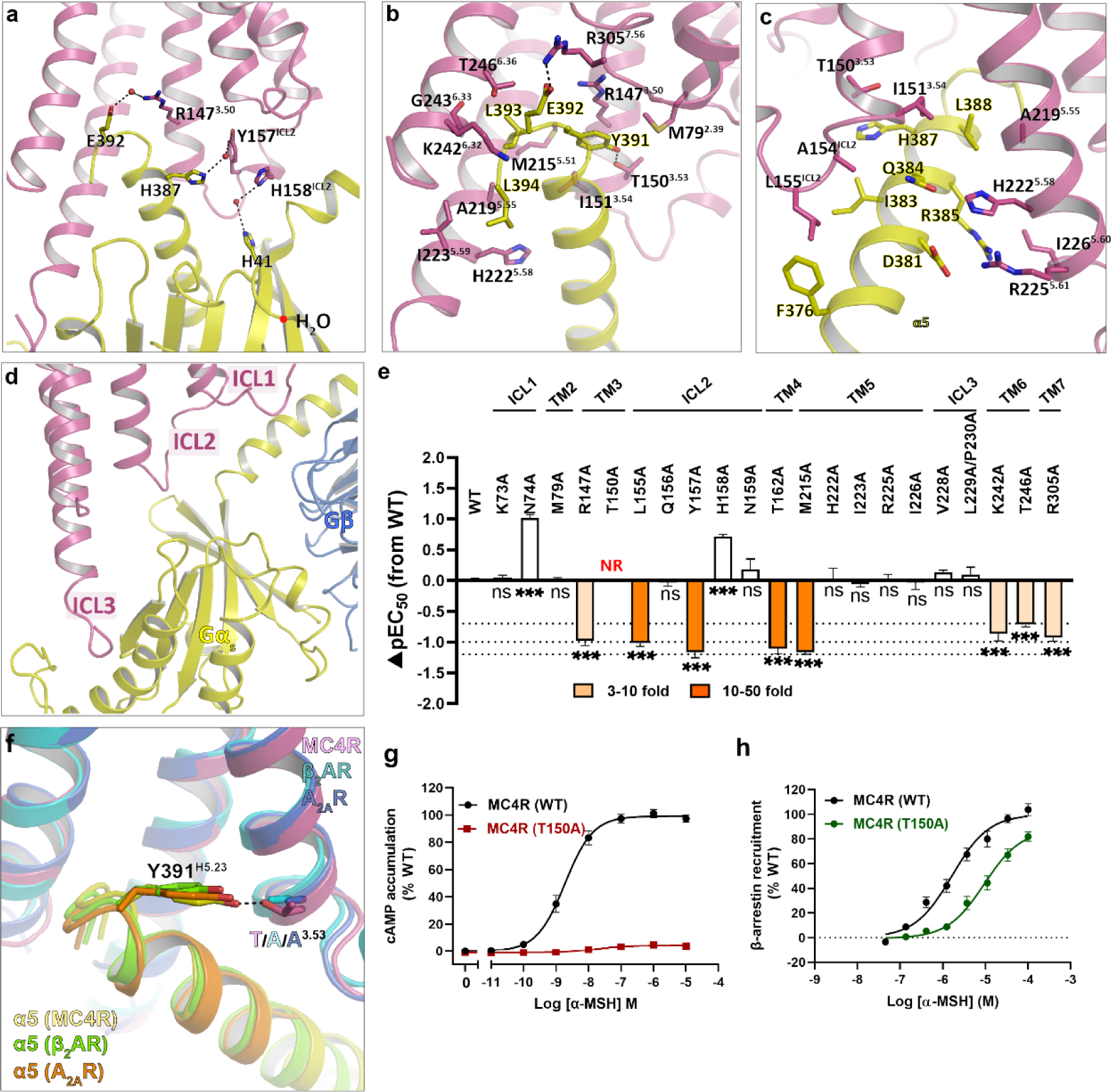
MC4R-Gs protein coupling. **a** Water molecules in the MC4R-Gs interface. Hydrogen bonds are shown as dashed lines. **b-c** Detailed interaction between MC4R and Gs-α5 helix. **d** Detailed interaction between three intracellular loops of MC4R and Gαs. **e** α-MSH-induced cAMP accumulation assays of the Gs binding site in MC4R. Bars represent differences in calculated α-MSH potency [pEC50] for each mutant relative to the wild-type receptor (WT). Data are colored according to the extent of effect. \*\**P* > 0.01, \**P* < 0.01, \*\**P* < 0.001 and \*\*\**P* < 0.0001 (one-way ANOVA followed by Dunnett’s multiple comparisons test, compared with the response of WT). **f** T^3.53^ of MC4R participates in the recognition of Y391 in Gs-α5 but not the corresponding A^3.53^ of β2AR and A2AR. Plum cartoon, MC4R structure; cyan cartoon, β2AR structure; blue cartoon, A2AR structure. MC4R–Gs, plum (MC4R) and yellow (α5); β2AR–Gs (PDB: 3SN6), cyan (β2AR) and green (α5); A2AR–Gs (PDB: 6GDG), blue (A2AR) and orange (α5). **g** α-MSH–induced cAMP accumulation assays of WT and T150^3.53^A mutant of MC4R. **h** α-MSH–induced β-arrestin2 recruitment assays of WT and T150^3.53^A mutant of MC4R. See Supplementary Table S4 and S14 for detailed statistical evaluation and receptor expression levels.

Of note, we observed multiple well-defined water molecules participating in the Gαs engagement to MC4R. To our knowledge, this is the first description of water molecule involvement in G protein coupling among class A GPCRs. It appears that water molecules connect three pairs of residues in the interface, E392^G.H5.24^ to R147^3.50^, H387^G.H5.19^ to Y157^ICL2^ and H41^G.S1.02^ to H158^ICL2^ (Fig. 7a). E392^G.H5.24^ also forms a salt bridge with R305^7.56^ in the TM7-H8 hinge, similar to other class A GPCRs (Fig. 7b). We individually mutated these residues in the Gs pocket, and assessed their effects on expression level and ability to stimulate a cAMP response. The results showed that R147^3.50^A, Y157^ICL2^A and R305^7.56^A reduced ligand potency 5-20-fold, pointing to crucial roles of these residues in Gs signaling (Fig. 7e; details are given in Supplementary information, Table S4). Though all the ICLs are involved in the Gs coupling, our mutagenesis study suggested that ICL2 played a more critical role compared to ICL1 and ICL3, supported by the fact substitutions of L155^ICL2^A and Y157^ICL2^A decreased α-MSH-induced receptor activation 10-20-fold whereas the mutations of residues in ICL1or ICL3 barely affected the ligand activity (Fig. 7e; details are given in Supplementary information, Table S4).

### Naturally occurring mutations in MC4R

To date, more than 200 genetic variants have been discovered in the MC4R gene ^57^. Human genetic studies show that loss-of-function (LoF) mutations in the MC4R gene are the most common monogenic cause of obesity. However, recent genetic association studies of approximately 0.5 million samples from the UK Biobank indicate that a subset of MC4R variants can cause gain-of-function (GoF) phenotypes, such as G231^ICL3^S, F201^5.47^L, I251^6.41^L and L304^7.55^F. Especially, GoF genetic variants in MC4R exhibited a signal bias for β-arrestin recruitment over canonical Gαs-mediated cAMP production and may have implications in obesity prevention.^8^ Therefore, structural determination of the activated MC4R-Gs complex offers a great opportunity to examine the molecular mechanism of functionally defective or enhanced variants.

Mutation of G231^ICL3^ to serine could potentially form a hydrogen bond with the spatially neighboring residue T350^G.h4S6.03^ of Gα resulting in enhanced Gαs-MC4R interaction, thus perfectly explaining the increased Gαs-mediated cAMP production by G231^ICL3^S mutation.^8^ However, another natural mutation at this position G231^ICL3^V that lacks the hydrogen bond with T350^G.h4S6.03^ did not affect Gs signaling.^8^ Interestingly, G231^ICL3^V/S variants in MC4R did not affect β-arrestin recruitment, indicating that G231^ICL3^ may not participate in β-arrestin signaling. In close proximity to the critical microswitch “toggle switch” and P/MIF motif, F201^5.47^L and I251^6.41^L variants may stabilize the receptor in the active state, leading to higher efficacies. At the cytoplasmic side, the hinge betweenTM7-H8 connects these two helices to TM1-ICL1-TM2, and plays important roles in G protein coupling and β-arrestin recruitment. Mutation of L304^7.55^ to phenylalanine with a larger side chain, which potentially increases the conformational rigidity of TM7-TM8-TM1-TM2 region by forming more hydrophobic interactions with the surrounding residues including L309^8.50^, R310^8.51^ and F313^8.54^, increased cAMP production and β-arrestin recruitment.^8^

A classification system was proposed to sort obesity-associated MC4R mutations.^58^ Many mutations that were functionally characterized as primarily defective in trafficking (class II), or with no apparent defect (class V) are actually involved in G-protein coupling according to our structural information and functional experiments. For example, T150^3.53^, Y157^ICL2^ and R305^7.56^ (class II) as well as Q156^ICL2^ and T162^4.38^ (class V) were found at the G-protein interface, and mutagenesis data show that these residues are crucial for Gs coupling (Fig.7d). Therefore, these mutants should also be included in class IV (defective in coupling to Gαs). Additionally, functional results indicate that I317^8.58^ mutants (class IV) did not affect G protein signaling,^8^ consistent with our structural observation that I317^8.58^ does not obviously interact with the G protein. Thus, it should not be classified as class IV (defective coupling to Gαs) mutants. ^52^ Naturally occurring mutants are closely related to obesity, and the detailed functional studies and accurate classifications are of great significance for practicing precision medicine.

## DISCUSSION

Delineating the structural basis for ligand recognition and G protein coupling at MC4R will aid in the rational design of drugs with high selectivity and reduced side effects. It will also facilitate our knowledge about naturally occurring MC4R mutants in the context of disease treatments. Here, we reported high-resolution cryo-EM structures of MC4R signaling complexes bound to four agonists with distinct chemical and functional features, including three non-selective peptidic agonists, *i.e.* endogenous peptide α-MSH, linear peptide afamelanotide and cyclic peptide bremelanotide, and one MC4R-selective small-molecule compound THIQ. Both peptide agonists and THIQ bind to the center of the wild-open binding pocket in the extracellular half of the MC4R 7TM bundle. The “toggle twin switch” L133^3.36^ and W258^6.48^, packed against the ligand phenyl ring, sense ligand binding and determine the functional nature of ligands. These structures together with mutagenesis studies revealed a common recognition mode of peptide agonists of the MCR family as well as critical determinants including TM3 residue I129^3.32^, ECL2 residue S188 and TM6 residue Y268^6.58^ responsible for THIQ selectivity. Furthermore, the structures also disclosed the MC4R-Gs engaging interface, providing the molecular basis for G protein-coupling specificity. In addition, all three ICLs of MC4R are involved in direct interactions with the Gs trimer, which may contribute to the high basal activity of MC4R. Together, our results offer unprecedented structural insights into the pharmacological and signaling features of MC4R and deepen our understanding of how naturally occurring mutants of MC4R modify the receptor activity. Our findings will serve as a structural template for rational drug design targeting the leptin-melanocortin pathway and facilitate the discovery of novel therapeutics against obesity. Moreover, these observed characteristics in the MC4R structures, including structural arrangement of the 7TM bundle, ligand-binding mode, agonist-induced receptor activation, and the manner of G protein coupling, could also be shared by other MCR family members.

## METHODS

### Cloning and insect cell expression

The wild-type (WT) human melanocortin-4 receptor (UniProt ID: P32245) gene was cloned into a modified pFastBac1 vector with hemagglutinin (HA) signal sequence at the N terminus and a PreScission protease site followed by a Flag tag. LgBiT and a double MBP tag were fused to the C terminus with a 3C protease cleavage site between them. A dominant-negative human Gαs (DNGαs) was generated by site-directed mutagenesis as previously described to stabilize the interaction with the βγ subunits ^31,59^ and cloned into pFastaBac1 vector. Gβ1 was fused with an N-terminal HiBit and 10 × His tag, together with Gγ2 were cloned into pFastBac dual vector.

The WT MC4R, DNGαs and Gβ1γ2 constructs were co-expressed in *Spodoptera frugiperda* (*Sf*9) insect cells using the Bac-to-Bac Baculovirus Expression System (Invitrogen). Cells were infected at a density of 2.4 × 10^6^ cells per mL and then co-infected with three separate viruses at a ratio of 1:1:1 for MC4R, DNGαs and Gβ1γ2. Cells were collected after 48 h post-infection and stored at −80°C until use.

### Expression and purification of Nb35

Nb35 with a C-terminal 6 × His-tag was expressed in the periplasm of *Escherichia. Coli* WK6 cells. The cells containing the recombinant plasmid were cultured in TB media supplemented with 0.1% glucose, 1 mM MgCl2 and 50μg/mL kanamycin at 37°C until OD600 reached 0.6. Then the cultures were induced by 1 mM IPTG and grown at 18°C for 24 h. The cells were harvested by centrifugation at 4,000 *g* for 15 min, and subsequently lysed. The protein was purified by nickel affinity chromatography as previously described ^34^. Eluted protein was concentrated using a 10 kDa molecular weight cut-off concentrator (Millipore) and loaded onto a HiLoad 16/600 Superdex 75 column (GE Healthcare) with running buffer containing 20 mM HEPES pH 7.5 and 100 mM NaCl. The monomeric fractions were pooled. Purified Nb35 was finally flash frozen in liquid nitrogen and stored at −80°C.

### Purification of MC4R**–**Gs**–**Nb35 complexes

The cell pellets were thawed on ice and lysed in a buffer containing 20 mM HEPES, pH 7.5, 100 mM NaCl and 2 mM MgCl2 supplemented with EDTA-free protease inhibitor cocktail (Bimake) by dounce homogenization. The complex formation was initiated by addition of 10 μg/mL Nb35, 25 mU/mL apyrase (NEB) and adequate agonist (100 μM α-MSH, 100 μM afamelanotide, 20 μM bremelanotide; China Peptides) or 10 μM THIQ (TopScience). The cell lysate was subsequently incubated for 1 h at room temperature (RT) and then solubilized by 0.5% (w/v) lauryl maltose neopentyl glycol (LMNG, Anatrace) and 0.1% (w/v) cholesterol hemisuccinate (CHS, Anatrace) for 2 h at 4°C. After centrifugation at 30,000 *g* for 30 min, the supernatant was isolated and incubated with amylose resin (NEB) for 2 h at 4°C. Then the resin was collected by centrifugation at 600 *g* for 10 min and loaded into a gravity flow column (Beyotime), and washed with five column volumes (CVs) of buffer containing 20 mM HEPES, pH 7.5, 100 mM NaCl, 2 mM MgCl2, agonist (100 μM α-MSH, 100 μM afamelanotide, 20 μM bremelanotide or 10 μM THIQ), 0.01% (w/v) LMNG and 0.005% (w/v) CHS, eluted with 15 CVs of buffer containing 20 mM HEPES, pH 7.5, 100 mM NaCl, 2 mM MgCl2, agonist (100 μM α-MSH, 10 μM afamelanotide, 20 μM bremelanotide or 10 μM THIQ), 0.01% (w/v) LMNG, 0.005% (w/v) CHS and 10 mM maltose. The elution was collected and incubated with 3C protease for 1 h at RT. Then the elution was concentrated with a 100 kDa cut-off concentrator (Millipore). Concentrated complex was loaded onto a Superose 6 increase 10/300 GL column (GE Healthcare) with running buffer containing 20 mM HEPES, pH 7.5, 100 mM NaCl, 2 mM MgCl2, agonist (100 μM α-MSH, 10 μM afamelanotide, 20 μM bremelanotide or 10 μM THIQ), 0.00075% (w/v) LMNG, 0.0002% (w/v) CHS and 0.00025% (w/v) GDN (Anatrace). The fractions for the monomeric complex were collected and concentrated for electron microscopy experiments.

### Cryo-EM grid preparation and data collection

For cryo-EM grid preparation, 3 μL of the purified α-MSH–, afamelanotide–, bremelanotide– or THIQ–bound MC4R–Gs complexes at ∼ 4 mg/mL were applied onto glow-discharged holey carbon grids (Quantifoil R1.2/1.3). Excess samples were blotted away and the grids were vitrified by plunging into liquid ethane using a Vitrobot Mark IV (ThermoFischer Scientific).

Cryo-EM imaging was performed on a Titan Krios at 300 kV in the Center of Cryo-Electron Microscopy, Zhejiang University (Hangzhou, China). Micrographs were recorded using a Gatan K2 Summit detector in counting mode with a pixel size of 1.014 Å using the SerialEM software. Movies were obtained at a dose rate of about 7.8 e/Å^2^/s with a defocus ranging from −0.5 to −2.0 μm. The exposure time was 8 s and 40 frames were recorded per micrograph. A total of 3,321, 3,521, 3,018 and 5,731 movies were collected for the α-MSH–, afamelanotide–, bremelanotide– and THIQ–bound MC4R– Gs complexes, respectively.

### Cryo-EM data processing

Image stacks were aligned using MotionCor 2.1.^60^ Contrast transfer function (CTF) parameters were estimated by Gctf v1.18.^61^ The following data processing was performed using RELION-3.0-beta2.^62^

For the α-MSH–bound MC4R–Gs complex, automated particle selection using Gaussian blob detection produced 2,170,482 particles. The particles were subjected to reference-free 2D classification to discard fuzzy particles, resulting in 680,962 particles for further processing. The map of GPBAR–Gs complex (EMD-30344)^56^ low-pass filtered to 60 Å was used as the reference map for 3D classification, generating one well-defined subset with 167,001 particles. Further 3D classifications focusing the alignment on the complex, produced three good subsets accounting for 147,565 particles, which were subsequently subjected to 3D refinement, CTF refinement and Bayesian polishing. The final refinement generated a map with an indicated global resolution of 3.0 Å at a Fourier shell correlation of 0.143.

For the afamelanotide–bound complex, 2,179,940 particles generated from the automated particle picking were subjected to 2D classification, producing 1,137,998 particles for 3D classification. The map of the α-MSH–bound complex was used as the reference for initial 3D classification, producing one good subset with 470,528 particles. Further 2 rounds of 3D classifications focusing the alignment on the MC4R–Gs complex or the MC4R receptor produced two high-quality subsets accounting for 217,491 particles, which were subsequently subjected to 3D refinement, CTF refinement and Bayesian polishing. The final refinement generated a map with an indicated global resolution of 3.0 Å.

For the bremelanotide–bound complex, automated particle picking yielded 2,259,228 particles, which were subjected to 2D classification. The well-defined classes were selected, producing 902,777 particles for further processing. The density map of α-MSH-bound complex low-pass filtered to 40 Å was used as the reference map for 3D classification, producing two good subsets with 274,222 particles. The particles were subsequently subjected to 3D refinement, CTF refinement and Bayesian polishing. The final refinement generated a map with an indicated global resolution of 3.1 Å.

For the THIQ–bound complex, automated particle picking yielded 2,905,757 particles, which were subjected to 2D classification. The well-defined classes were selected, producing 1,148,249 particles for further processing. The density map of α-MSH-bound complex low-pass filtered to 40 Å was used as the reference map for 3D classification, producing two good subsets with 269,832 particles. The particles were subsequently subjected to 3D refinement, CTF refinement and Bayesian polishing. The final refinement generated a map with an indicated global resolution of 2.9 Å.

Local resolution was determined using the Bsoft package with half maps as input maps.^63^

### Model building, refinement and validation

The initial homology model of the active state MC4R was generated from GPCRdb.^42^ The parathyroid hormone receptor-1–Gs–Nb35 complex (PDB 6NBF)^64^ was to generate the initial models of Gs and Nb35. Small molecular ligand coordinates and geometry restraints were generated using phenix.elbow. Models were docked into the EM density map using UCSF Chimera (https://www.cgl.ucsf.edu/chimera/). This starting model was then subjected to iterative rounds of manual adjustment and automated refinement in Coot^65^ and Phenix^66^, respectively.

The final refinement statistics were validated using the module ‘comprehensive validation (cryo-EM)’ in PHENIX. To monitor the potential over-fitting in model building, FSCwork and FSCtest were determined by refining the ‘shaken’ models against unfiltered half-map-1 and calculating the FSC of the refined models against unfiltered half-map-1 and half-map-2. Structural figures were prepared in Chimera (https://www.cgl.ucsf.edu/chimera), ChimeraX^67, 68^ and PyMOL (https://pymol.org/2/). The final refinement statistics are provided in Supplementary Table S2.

### cAMP accumulation assay

Agonist (α-MSH, afamelanotide, bremelanotide or THIQ) stimulated cAMP accumulation was measured by a GloSensor™ cAMP assay kit (Promega). Briefly, HEK293T cells transfected with WT or mutant MC4R and the pGloSensor™-22F plasmid were seeded onto 384-well culture plates at a density of 4 × 10^3^ cells per well and incubated for 24 h at 37°C in 5% CO2. Then the culture medium was removed and the equilibration medium containing 4% (v/v) dilution of the GloSensor™ cAMP reagent stock solution was added to each well. To analyze the effect of Ca^2+^ on cAMP signaling, the equilibration medium was further supplemented with different concentrations of EGTA (0 mM, 1 mM, 5 mM, 10 mM and 20 mM). To obtain the concentration-response curves, serially diluted agonists were added to each well to stimulate the cells.

Luminance signal was measured using 0.5 s intervals after ligand addition (TECAN, 25°C). Concentration-responses were generated from the peak response. cAMP accumulation was analyzed by a standard dose-response curve using GraphPad Prism 8.0 (GraphPad Software). EC50 and pEC50 ± SEM were calculated using nonlinear regression (curve fit). Data are means ± SEM from at least three independent experiments performed in technical triplicates.

### Receptor expression

Plasmids corresponding to WT and mutant MC4R, or pcDNA3.1 (+) empty vector, were transfected as described above. After transfection, cells were re-seeded onto poly-D-lysine treated 96-well plates at a density of 3 × 10^4^ cells per well. Twenty-four hours later, cells were washed with PBS and fixed with 10% formaldehyde for 10 min followed by three times washing with PBS. Following fixation, cells were blocked with blocking buffer (1% BSA in PBS) for 1 h at RT. Afterward, plates were incubated with a 1:20,000 dilution of anti-FLAG M2 HRP-conjugated monoclonal antibody (Sigma-Aldrich) in blocking buffer for another 1 h at RT. After careful washing, 80 μL/well diluent SuperSignal Elisa Femto Maximum Sensitivity Substrate (ThermoFisher Scientific) was added. Luminance signal was measured using 1 s intervals. Data are means ± SEM from at least three independent experiments performed in technical triplicates.

### Radioligand binding assay

Radioligand binding assays for WT and mutant MC4R were performed in transiently transfected HEK293 cells. Briefly, cells were seeded at a density of 3 × 10^4^ cells/well onto 96-well culture plates and incubated overnight at 37°C in 5% CO2, and competitive radioligand binding was assessed 24 h thereafter. For homogeneous binding, the cells were incubated in binding buffer (DMEM supplemented with 25 mM HEPES and 0.1% BSA) with a constant concentration of ^125^I-α-MSH, (40 pM, PerkinElmer) and different concentrations of unlabeled peptides [α-MSH (47.68 pM to 100 µM); afamelanotide (1 pM to 10 µM); bremelanotide (1 pM to 10 µM)] at RT for 3 h. Then plates were washed three times with ice-cold PBS and lysed by 50 μl lysis buffer (PBS supplemented with 20 mM Tris-HCl, pH 7.4 and 1% Triton X-100). The plates were subsequently counted for radioactivity (counts per minute, CPM) with a MicroBeta^2^ plate counter (PerkinElmer) using a scintillation cocktail (OptiPhase SuperMix, PerkinElmer).

### β-arrestin 2 recruitment assay

β-arrestin 2 recruitment was measured in HEK293T cells using a bioluminescence resonance energy transfer (BRET) assay. The cells were seeded onto poly-D-lysine coated 96-well plates at a density of 3.5 × 10^4^ cells/well and grown overnight before transfection. Prior to BRET experiments, cells were transiently transfected with MC4R-Rluc8 and β-arrestin 2-Venus at a ratio of 1:6 for 24 h and rinsed twice with HBSS. The cells were then incubated with BRET buffer (HBSS supplemented with 10 mM HEPES and 0.1% BSA, pH 7.4) for 30 min at 37°C. After incubation with 5 μM coelenterazine-H (Yeasen Biotech) for 5 min, the baseline BRET signals were read immediately at 470 nm and 535 nm for 10 cycles using an EnVision^®^ multimode plate reader (PerkinElmer). Following the agonist addition, BRET was measured every 1 min for another 50 cycles. Data are presented as BRET ratio, calculated as the ratio of Venus to Rluc8 signals and subtracted the vehicle value. Concentration-response profiles were obtained from area-under-the-curve values of elicited responses.

### Molecular docking

The active MC4R model was generated from the high-resolution structure of α-MSH– MC4R–Gs complex for molecular docking. Three representative analogs of small molecule THIQ were docked into the active MC4R model using the “RosettaLigand” application in Rosetta 2019.35 version.^69, 70^ Conformers of the three ligands were generated by the Open Babel program (The Open Babel Package, version 2.3.1 http://openbabel.org).^71^ Then the ligand docking poses were searched from the pocket of MC4R with a box size of 15 Å controlled by the RosettaScrpits. A total of 1,000 models were generated for the docking trial. The top 10 docking poses of each ligand were selected based on the lowest binding energy (interface_delta_X) and viewed by PyMol. The most similar binding pose as THIQ of each ligand was selected and displayed in Supplementary Figure S6.

### Statistical analysis

All functional study data were analyzed using Prism 8 (GraphPad) and presented as means ± S.E.M. from at least three independent experiments. Concentration-response curves were evaluated with a standard dose-response curve. Statistical differences were determined by two-sided, one-way ANOVA with Dunnett’s multiple comparisons test.

### Data availability

The atomic coordinates and the electron microscopy maps of the α-MSH–, afamelanotide–, bremelanotide– and THIQ–bound MC4R–Gs complexes have been deposited in the Protein Data Bank (PDB) under accession number xxx, xxx, xxx, xxx and Electron Microscopy Data Bank (EMDB) under accession codes xxx, xxx, xxx, xxx respectively. All relevant data are included in the manuscript or Supplementary Information.

## Acknowledgements

The cryo-EM data were collected at the Center of Cryo-Electron Microscopy, Zhejiang University. This work was partially supported by grants from National Natural Science Foundation of China g 81922071 (Y.Z.), 81872915/82073904 (M.-W.W.) and 81773792/81973373 (D.Y.); the National Key Basic Research Program of China 2019YFA0508800 (Y.Z.), 2018YFA0507000 (M.-W.W.) and 2018ZX09711002–002–005 (D.Y.); Zhejiang Province Science Fund for Distinguished Young Scholars LR19H310001 (Y.Z.); Key R & D Projects of Zhejiang Province 2021C03039 (Y.Z.); National Science & Technology Major Project of China – Key New Drug Creation and Manufacturing Program 2018ZX09735–001 (M.-W.W.) and 2018ZX09711002–002–005 (D.Y.); and Novo Nordisk-CAS Research Fund NNCAS-2017–1-CC (D.Y.).

## string-namehor contributions

H.Z. designed the expression constructs, purified all the MC4R−Gs complexes, prepared the final samples for negative staining EM and cryo-EM data collection, and participated in manuscript preparation; H.Z., D.D.S. and C.M. performed specimen screening, cryo-EM grid preparation and data collection, map calculation; H.Z., L.C., Q.S. and C.M. performed structural analysis and figures preparation; H.Z., L.C., S.Y., W.F., A.D., S.X., Y. Zhou and D.Y. conducted mutation studies, ligand binding and signaling experiments; Y.Z. and H.Z. analyzed the data with the support of L.C., J.S., D.H.S., D.Y., H.E.X, T.H., T.Z. and M.-W.W.; Y.Z., M.-W.W. and T.Z. supervised the project and wrote the manuscript with inputs from all co-string-namehors.

---

**Supplementary information, Fig. S1.**
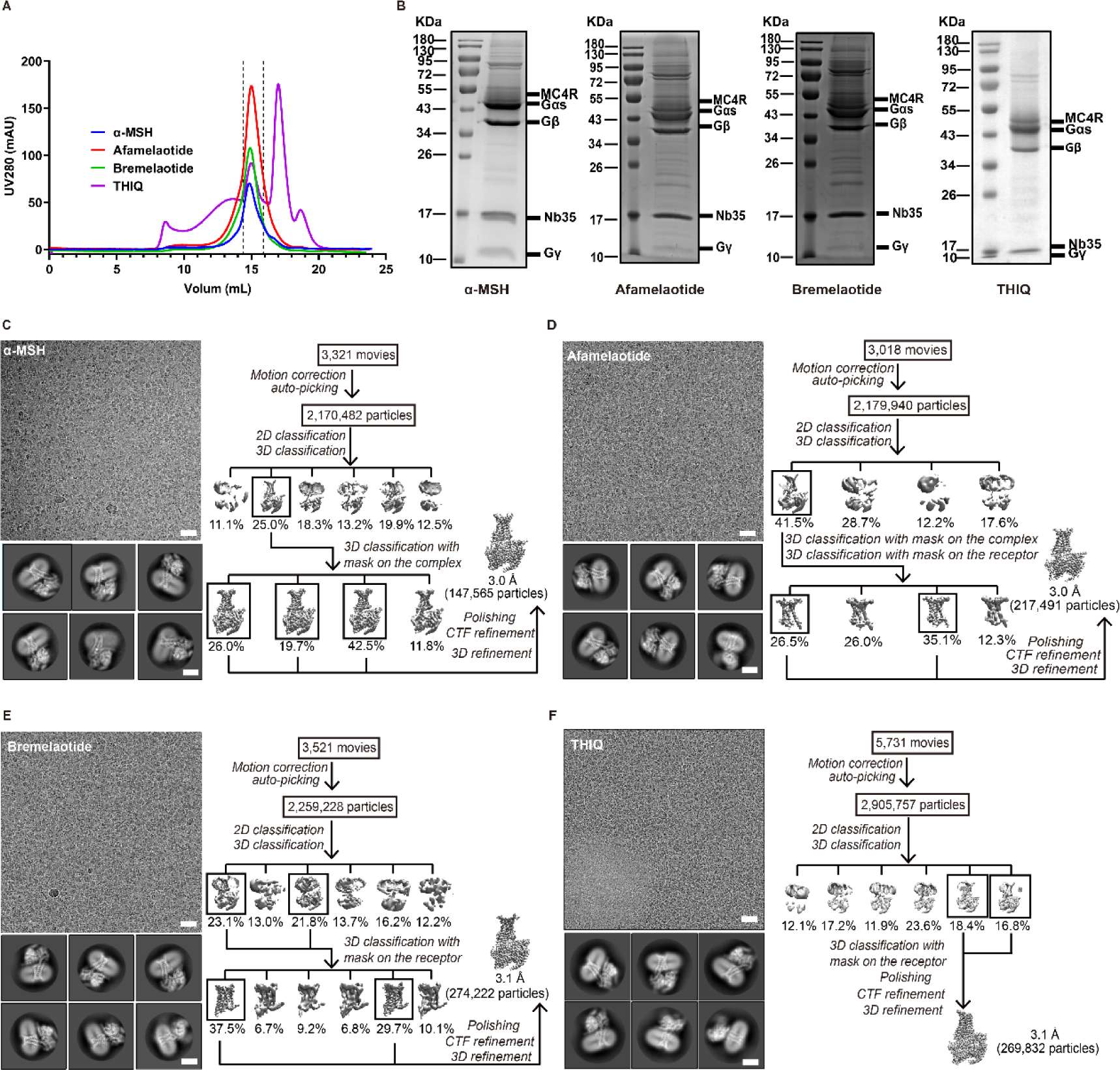
Purification and single-particle reconstruction of the agonists–bound MC4R–Gs complexes. a, Size exclusion chromatography (SEC) profile (a) and SDS-PAGE analysis (b) of α-MSH–, afamelanotide–, bremelanotide– and THIQ–bound MC4R–Gs complexes. Fractions between two dashed lines in the SEC profile were pooled and concentrated for cryo-EM analysis. c-f Cryo-EM micrographs (scale bar: 30 nm), 2D class averages (scale bar: 5 nm) and the flow chart of cryo-EM data processing for α-MSH– (c), afamelanotide– (d), bremelanotide– (e) and THIQ–bound (f) MC4R–Gs complexes.

**Supplementary information, Fig. S2.**
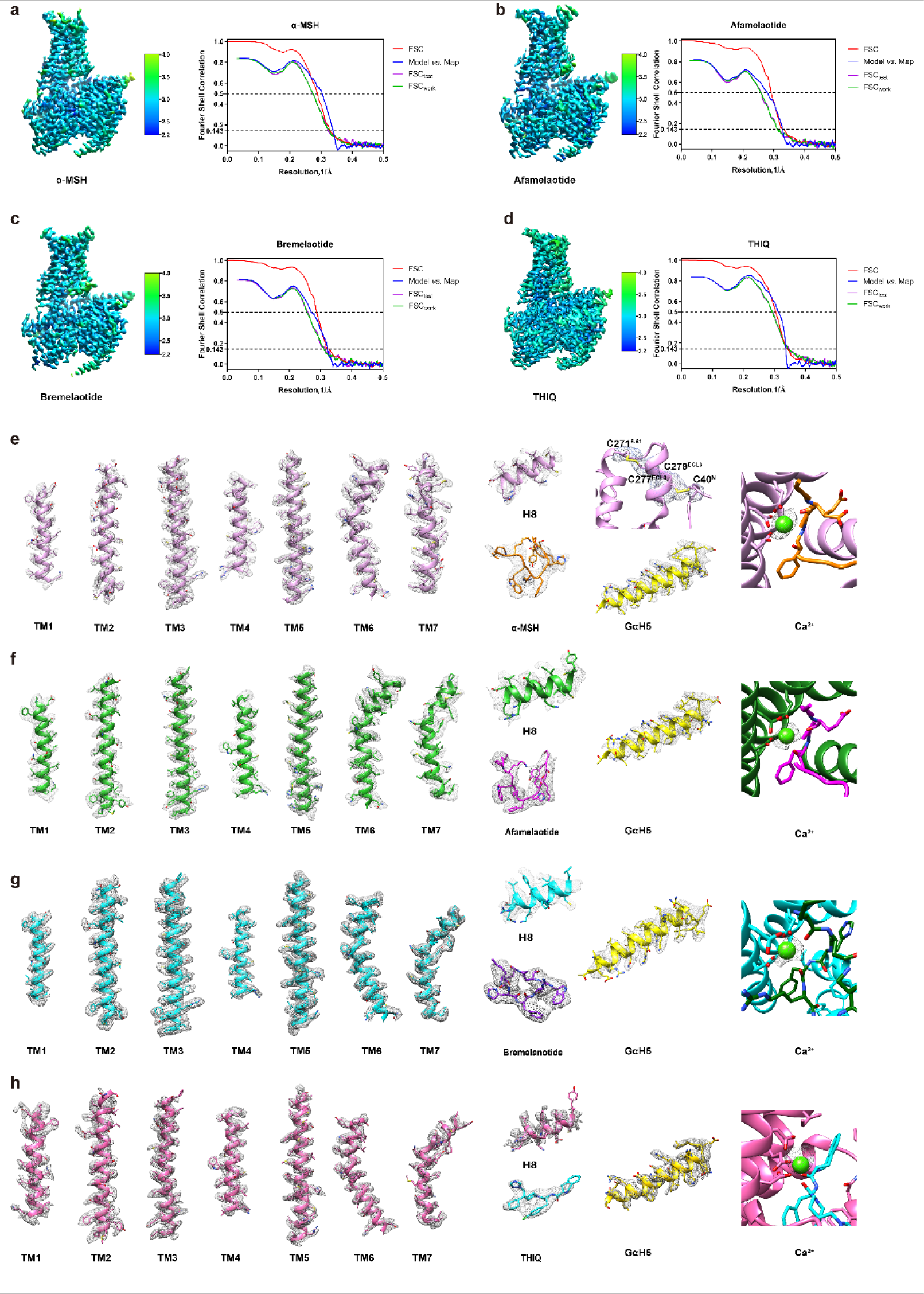
Overall resolution and cryo-EM density analysis of the MC4R-Gs complexes. a-d, Local resolution maps, gold-standard fourier shell correlation (FSC) curves, the model-vs-map, FSCwork and FSCtest validation curves of the α-MSH– (a), afamelanotide– (b), bremelanotide– (c) and THIQ–bound (d) MC4R–Gs complexes. e-h, EM density maps and models are shown for all seven-transmembrane helices, helix 8, agonists, Gαs α5-helix and Ca^2+^ of the α-MSH– (e), afamelanotide– (f), bremelanotide– (g) and THIQ–bound (h) MC4R–Gs complexes. α-MSH–bound MC4R, plum; afamelanotide–bound MC4R, green; bremelanotide–bound MC4R, cyan; THIQ– bound MC4R, pink; α-MSH, orange; afamelanotide, magenta; bremelanotide, purple; THIQ, cyan; Gαs-α5, yellow; Ca^2+^, green. Densities are shown at 0.020-0.045 contour level, except the disulfide bond between C40 and C279 at 0.01 contour level.

**Supplementary information, Fig. S3.**
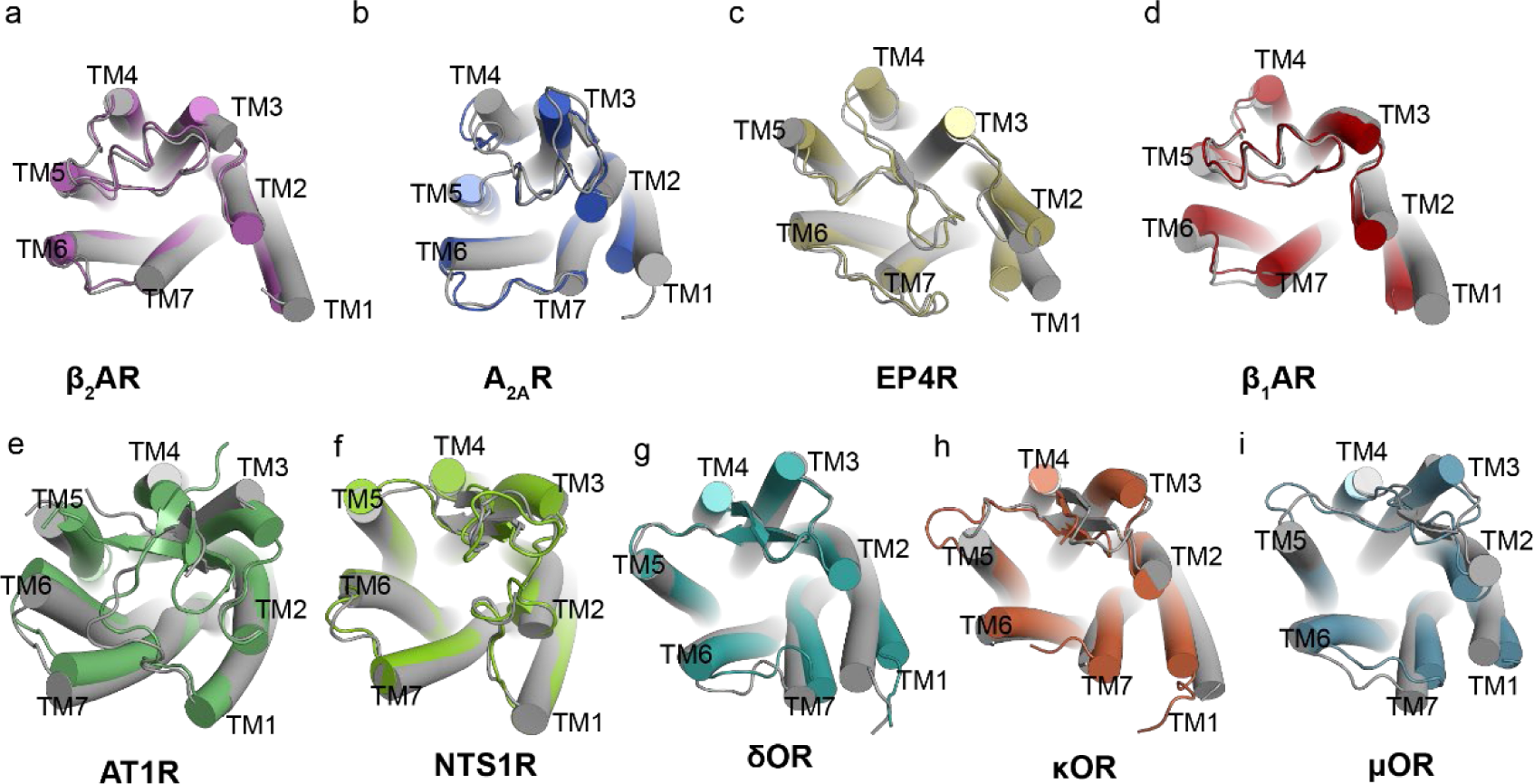
Structural comparison between active structures of class A Gs-coupled receptors and other peptide receptors. a-d, Superimposition of inactive and active structures of class A Gs-coupled receptors. Inactive and active β2AR, grey (PDB: 6PS6) and purple (PDB: 3SN6); inactive and active A2AR, grey (PDB: 6LPJ) and blue (PDB: 6GDG); inactive and active EP4, grey (PDB: 5YWY) and yellow (PDB: 7D7M); inactive and active β1AR, grey (PDB: 3ZPQ) and red (PDB: 7JJO). e-i, Superimposition of inactive and active structures of class A peptide receptors. Inactive and active AT1R, grey (PDB: 4YAY) and green (PDB: 6DO1); inactive and active NTSR1, grey (PDB: 4BUO) and yellow green (PDB: 6OS9); inactive and active δOR, grey (PDB: 4N6H) and turquoise (PDB: 6PT3); inactive and active κOR, grey (PDB: 4DJH) and orange (PDB: 6B73); inactive and active μOR, grey (PDB:4DKL) and steel blue (PDB:6DDE).

**Supplementary information, Fig. S4.**
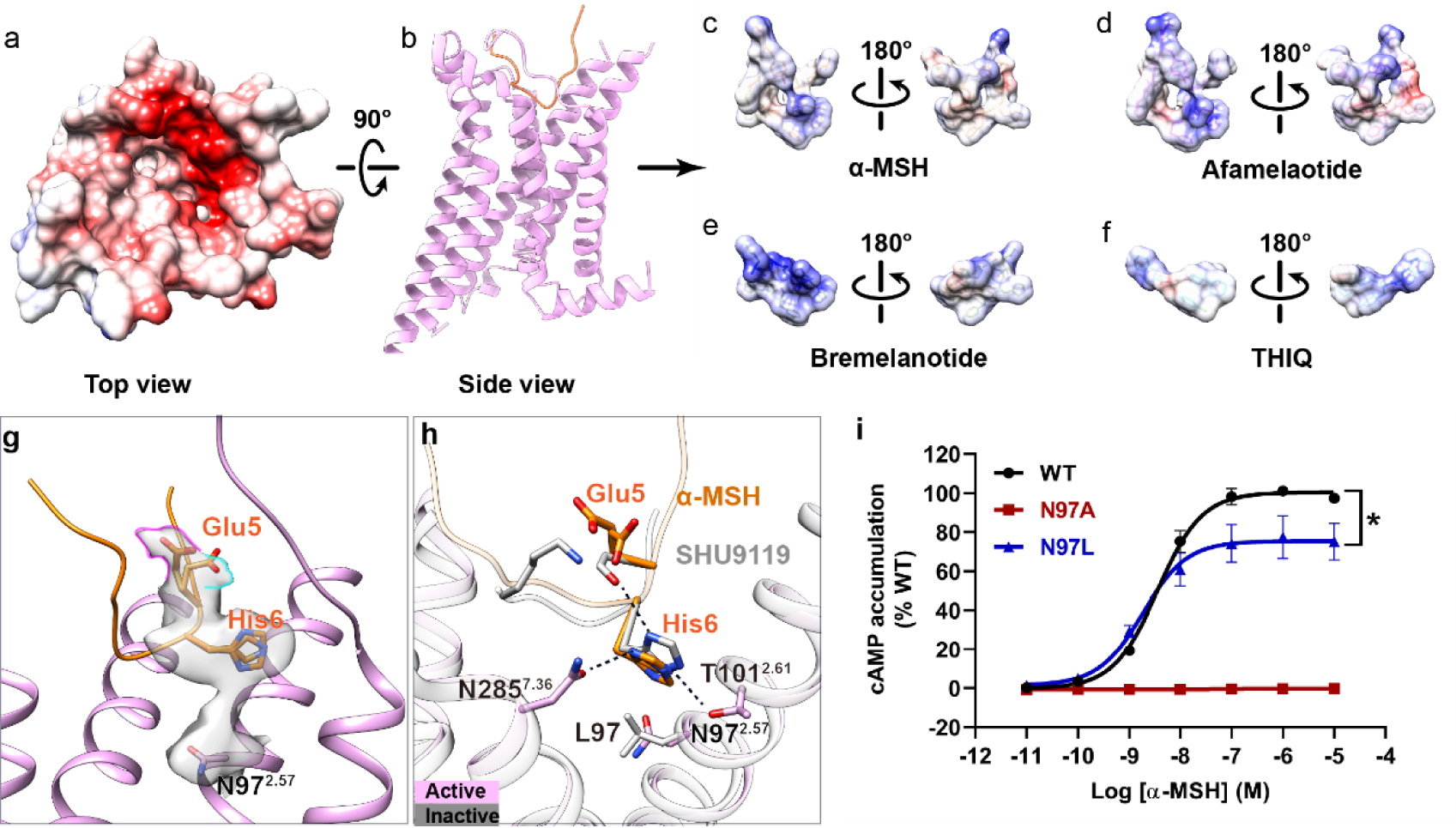
Ligand binding pocket of MC4R. a-f, The electrostatic potential analysis for the orthosteric pocket of MC4R (a-b) as well as α-MSH (c), afamelanotide (d), bremelanotide (e) and THIQ (f). g, Cryo-EM densities of Glu5 and His6 in α-MSH and N97^2.57^ in MC4R are shown at the 0.011 contour level. h, Superposition of the α-MSH (orange)–activated MC4R (plum) with the antagonist (SHU9119, orange)-bound and thermostabilized MC4R (N97^2.57^L) mutant (grey, PDB 6W25). i, α-MSH–induced cAMP accumulation assays of WT, N97A and N97L MC4Rs, \**P* < 0.05. (i) See Supplementary Table S4 and S5 for detailed statistical evaluation and receptor expression levels.

**Supplementary information, Fig. S5.**
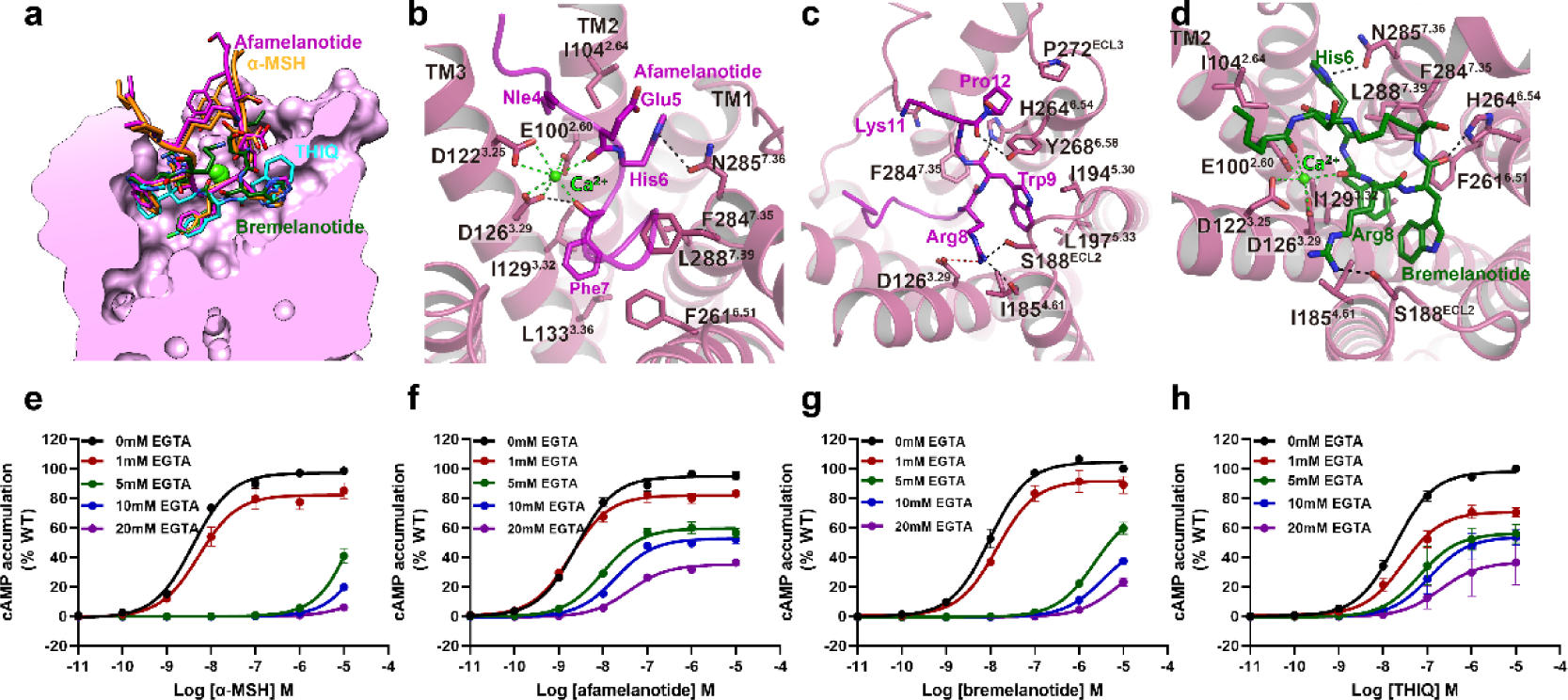
Binding sites of afamelanotide and bremelanotide in MC4R and effects of Ca^2+^ on agonist binding and agonist– induced signaling. a, Comparative binding poses of four different ligands in the binding pocket. b-d, Detailed interaction between afamelanotide (b-c) and bremelanotide (d) with MC4R. MC4R, plum; afamelanotide, magenta; bremelanotide, sea green; Ca^2+^, green. e-h, Effects of Ca^2+^ on α-MSH– (e), afamelanotide– (f), bremelanotide– (g) and THIQ– (h) induced cAMP accumulation.

**Supplementary information, Fig. S6.**
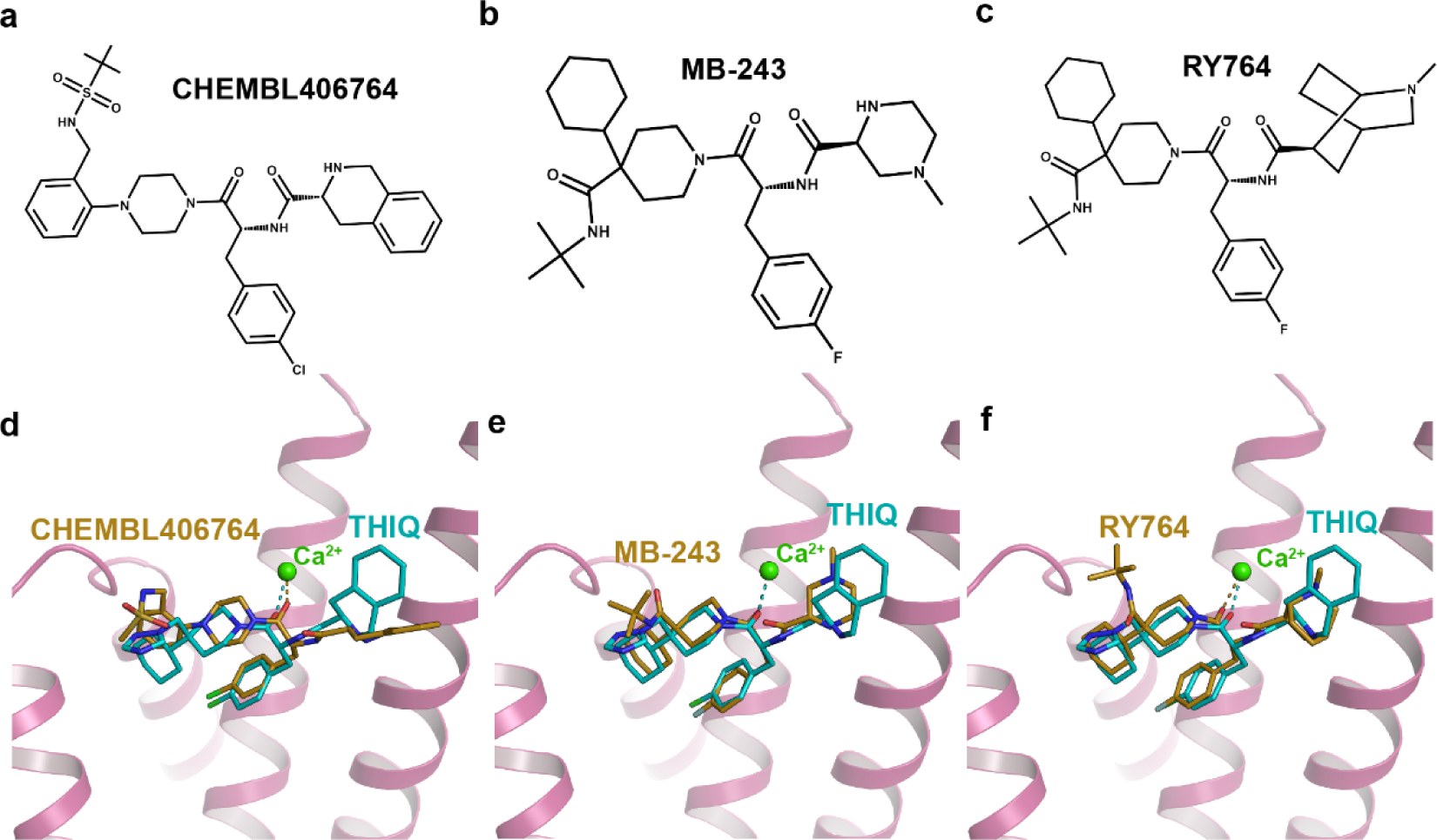
Molecular docking of MC4R agonists. a-c, Chemical structures of CHEMBL406764 (a), MB-243 (b) and RY764 (c); d-f, Docking of CHEMBL406764 (d), MB-243 (e) and RY764 (f) in the THIQ–bound MC4R structure. The rank 1 binding pose of each ligand was selected to display.

**Supplementary information, Fig. S7.**
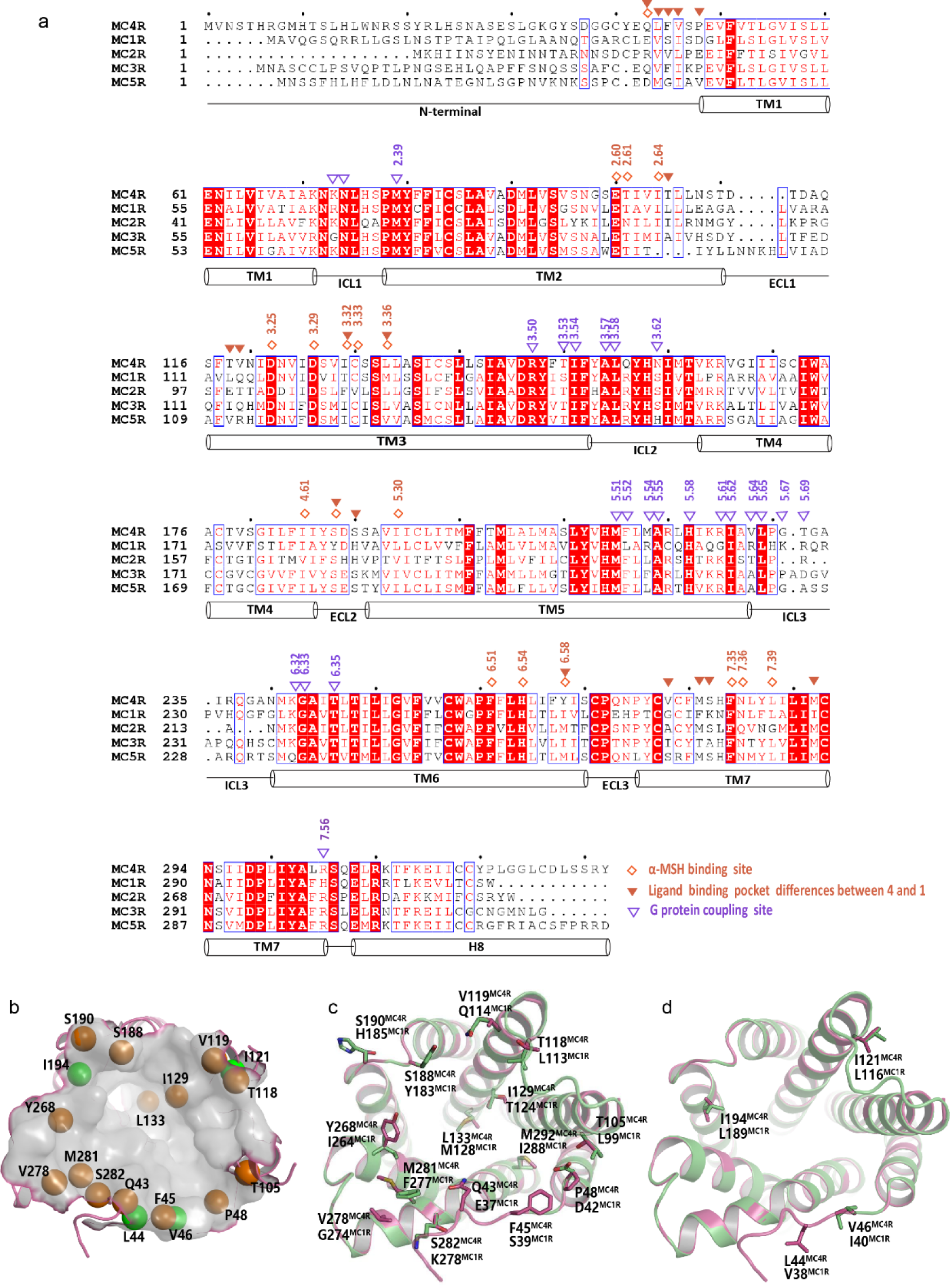
Sequence alignment of the MCR family members and the orthosteric pocket difference between MC4R and MC1R. a, Sequence alignment of the melanocortin family receptors. The sequence alignment was created using GPCRdb (http://www.gpcrdb. org). The residues of α-MSH binding sites, Gs coupling sites and different residues between MC4R and MC1R in respective ligand-binding pocket are highlighted with hollow diamond, hollow triangle and solid triangle symbols. Secondary structure elements are annotated underneath the sequences based on the structure of MC4R. b, The orthosteric pocket of MC4R. The pocket surface is displayed in grey and the residues in the ligand pocket of MC4R that are different from the corresponding residues of MC1R are shown as balls (the orange balls represent the residues displaying different chemical properties from the corresponding residues of MC1R, the green balls represent residues with similar chemical properties). c-d, Differentiated residues in the orthosteric pocket between the MC4R structure and the homology model of MC1R. c, Highlighting view of the residues with different chemical properties. d, Highlighting view of the residues with similar chemical properties.

**Supplementary information, Fig. S8.**
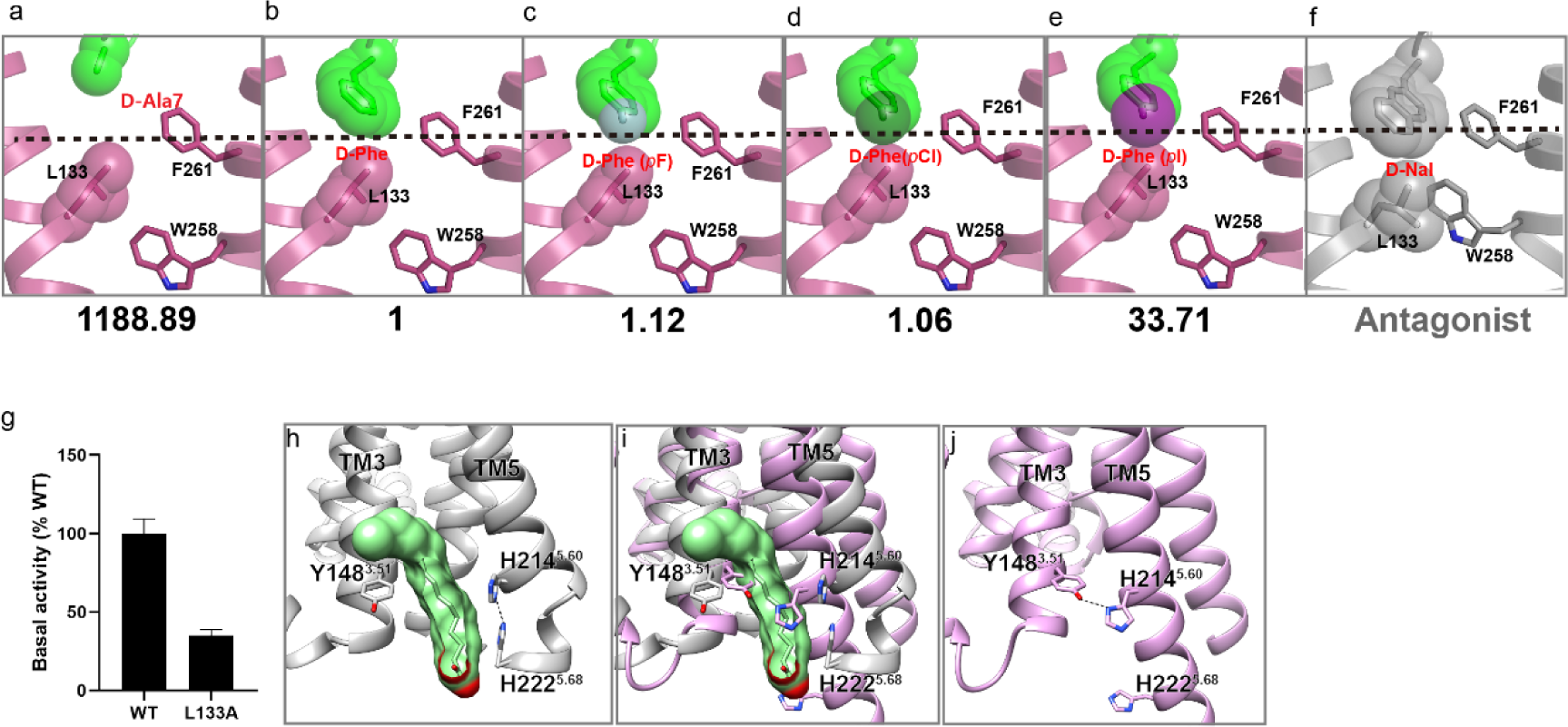
Relationships between chemical modification at the position of D-Phe residue and MC4R activation, and lipid in the inactive MC4R. a-f, Chemical replacement of D-Phe to D-Ala (a), D-Phe(*p*F) (c), D-Phe(*p*Cl) (d) and D-Phe(*p*I) (e) based on the active structure of bremelanotide-bound MC4R (b). The D-Nal modification is exhibited using the inactive structure of SHU9119-MC4R (f). The fold change in EC50 value of each ligand compared to [Nle4, D-Phe7]α-MSH-NH2 (b) is shown under the corresponding panel. Active MC4R, plum; inactive MC4R, grey. g, Basal activity of MC4R (wild type, WT) and MC4R (L133^3.36^A). h, Lipid between TM3 and TM5 in the inactive MC4R. i, Superimposition of inactive and active structures of MC4R. j, Hydrogen bond between Y148^3.51^ and H214^5.60^ in the inactive MC4R.

**Supplementary information, Fig. S9.**
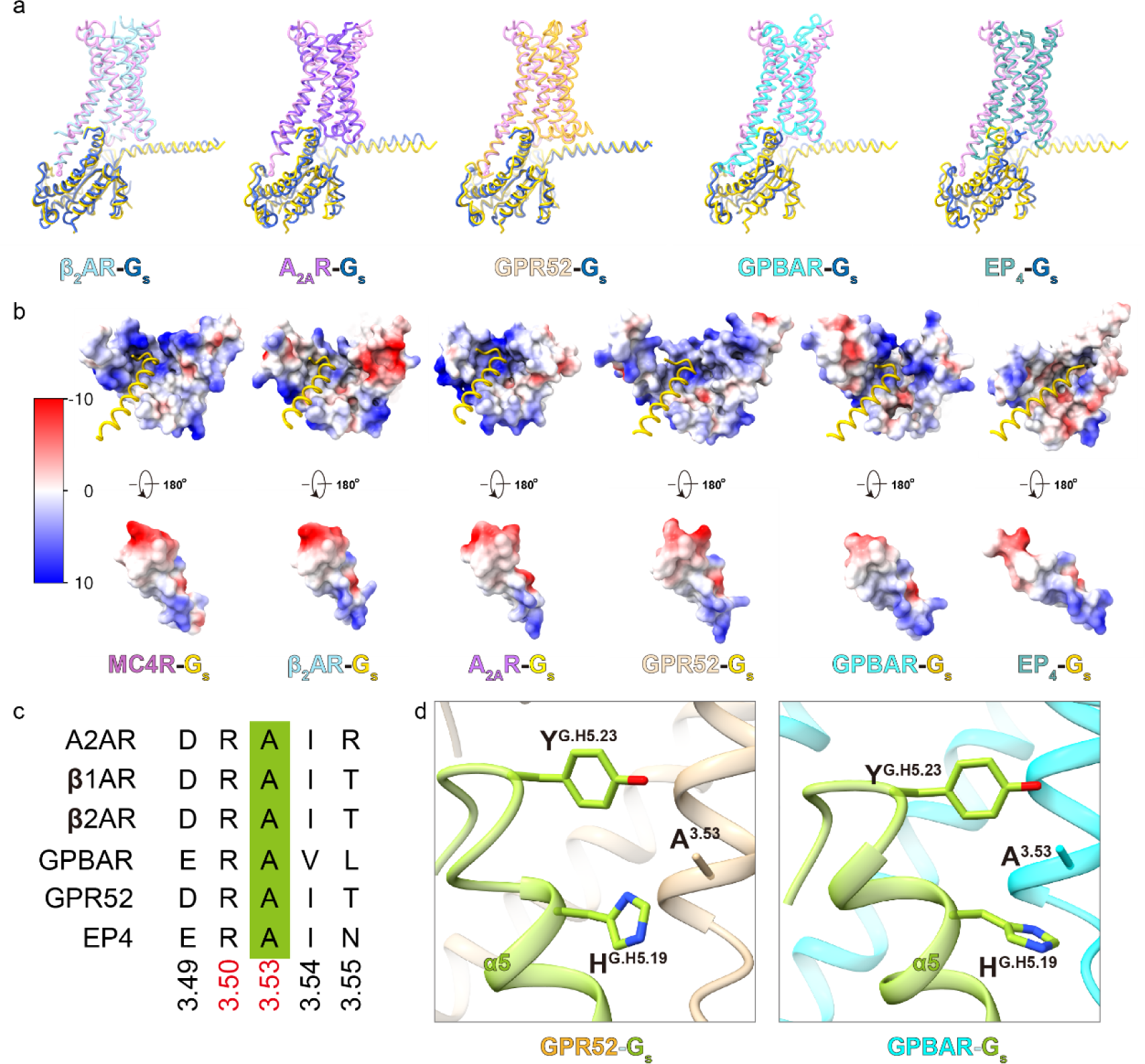
Interaction patterns between class A GPCRs and Gαs. a, Pairwise comparison of MC4R–Gαs and other class A GPCR–Gαs structures. MC4R, plum; MC4R–bound Gαs, yellow; other receptor-bound Gαs, blue; β2AR (PDB: 3SN6); sky blue; A2AR (PDB: 6GDG), purple; GPR52 (PDB: 6LI3); orange, GPBAR (PDB: 7CFM); cyan, EP4 (PDB: 7D7M), teal. b, The electrostatic potential analysis of the G protein interface for class A GPCR–Gs complexes. c, Residues on TM3 involved in G protein recognition among for Gs coupled class A receptors. The presentation was created using GPCRdb (http://www.gpcrdb.org). The conserved interaction residues of R^3.50^ and A^3.53^ are highlighted. d, Detailed interactions of receptor A^3.53^ with the α5 helix of G protein in the structures of GPR52–Gs and GPBAR–Gs.

